# Bacterial communities of *Aedes aegypti* mosquitoes differ between crop and midgut tissues

**DOI:** 10.1101/2022.08.31.506054

**Authors:** Luis E. Martinez Villegas, James Radl, George Dimopoulos, Sarah M. Short

## Abstract

Microbiota studies of *Aedes aegypti* and other mosquitoes generally focus on the bacterial communities found in adult female midguts. However, other compartments of the digestive tract maintain communities of bacteria which remain almost entirely unstudied. For example, the Dipteran crop stores nectar and other sugars, but few studies have looked at the microbiome of crops in mosquitoes, and only a single previous study has investigated the crop in *Ae. aegypti*. In this study, we used both culture-dependent and culture-independent methods to compare the bacterial communities in midguts and crops of laboratory-reared *Ae. aegypti*. Both methods revealed a trend towards higher abundance, but also higher variability, of bacteria in the midgut than the crop. When present, bacteria from the genus *Elizabethkingia* (family Weeksellaceae) dominated midgut bacterial communities. In crops, we found a higher diversity of bacteria, and these communities were generally dominated by acetic acid bacteria (family Acetobacteriaceae) from the genera *Tanticharoenia* and *Asaia*. These three taxa drove significant community structure differences between the tissues. We used FAPROTAX to predict the metabolic functions of these communities and found that crop bacterial communities were significantly more likely to contain bacteria capable of methanol oxidation and methylotrophy. Both the presence of acetic acid bacteria (which commonly catabolize sugar to produce acetic acid) and the functional profile that includes methanol oxidation (which is correlated with bacteria found with natural sources like nectar) may relate to the presence of sugar in the crop. A better understanding of what bacteria are present in the digestive tract of mosquitoes and how these communities assemble will inform how the microbiota impacts mosquito physiology and the full spectrum of functions provided by the microbiota. It may also facilitate better methods of engineering the mosquito microbiome for vector control or prevention of disease transmission.

**Author summary:** Bacteria inside mosquitoes’ guts have been found to have an impact on mosquito life history traits (such as longevity and fecundity) as well as their susceptibility to infection by human pathogens. Engineering these communities may provide an effective and safe way to control mosquitoes and reduce the impact of the pathogens they spread. In this work, we assayed the bacteria found in midgut and crop tissues of a medically important mosquito, *Aedes aegypti*. Our results show that these tissues harbor communities of bacteria that differ in composition and function and vary in abundance. Experiments like ours are important to better understand where bacteria are found in an insect’s body and how these communities assemble. This knowledge may help future researchers more successfully engineer bacterial communities in mosquitoes.

## Introduction

The yellow fever mosquito, *Aedes aegypti* (L.), transmits multiple human arboviral pathogens (including dengue virus, Zika virus, chikungunya virus, and yellow fever virus) that collectively cause more than one hundred million human infections each year and tens of thousands of deaths [1]. Vaccines exist for some of these arboviruses with varying efficacy [2,3]. However, many lack vaccines, and treatment is limited in all cases to supportive care. Because arboviral threats are often emerging and challenging to predict [4], control methods targeting mosquito vectors are often the primary mode to prevent transmission and reduce disease cases.

*Ae. aegypti* mosquitoes are found in close association with microbial communities throughout their lives. The most well-studied of these associations are the bacterial communities that inhabit the larval habitat and the larval and adult female digestive tract. As larvae, *Ae. aegypti* develop in stagnant water found primarily in man-made containers. Larvae ingest microbes routinely throughout development [5], and their gut microbiota is primarily composed of microbes orally acquired from the environment [6,7]. As adults, *Ae. aegypti* also host bacterial communities in their digestive tract. The sources of these bacteria are not clear. Bacteria can be transstadially transmitted from larvae to adults which may account for some of the bacteria found in adults [6,8,9], but it is also hypothesized that adults acquire bacteria when nectar feeding [10].

The microbial communities associated with adult *Ae. aegypti* mosquitoes have garnered much attention because of their sizeable impact on life history traits and susceptibility to infection by human pathogens (reviewed in [11]). For example, the microbiota of adults can impact susceptibility to dengue virus, blood meal digestion, and fecundity [12,13]. Additionally, experimentally introducing certain microbes to the digestive tract of adult female mosquitoes significantly impacts mosquito longevity and susceptibility to arboviruses [14–17].

The vast majority of investigations into the adult microbiota of mosquitoes has focused on the midgut portion of the digestive tract because the midgut is where the blood meal is stored during digestion and is the tissue through which blood borne pathogens invade [18,19]. However, microbes are also found in other compartments of the adult mosquito digestive tract, such as in the crop which is a nectar storage organ anterior to the midgut [20]. In mosquitoes and other adult hematophagous Diptera (e.g., Ceratopogonidae [21]; Simuliidae [22]), the crop is a ventral diverticulum of the esophagus (foregut) separated by a muscular valve [23]. Sensory cells signal the esophageal valve to direct meals rich in sugar (e.g. nectar or artificial sugar supplements) to the crop and meals low in sugar (e.g. blood meals) to the midgut [24–26]. Sugar-rich meals are stored in the crop until they are pumped by peristaltic contractions of the crop back into the digestive tract for digestion [27]. This allows the mosquito to retain sugars until they are needed for energy intensive activities, such as flight [28]. The crop is generally considered only a storage organ because its cuticular lining prevents most nutrients from being absorbed [23]. However, some carbohydrate digestion (e.g., sucrose catabolism) may occur here because some salivary enzymes are also retained with the meals in the crop [23,25]. The Dipteran crop is known to maintain stable populations of microbes (reviewed in [28,29]) which may also aid in the digestion of these sugars.

Little research to date has focused on the microbiota in the mosquito crop. A better understanding of the mosquito crop microbiota is of interest because (1) it could be contributing to the physiological impacts of the microbiota described above, either independently or in concert with the microbial communities in other regions of the gut, and (2) it is possible that microbes from the crop could influence formation of the midgut microbiota if they are passed from the crop to the midgut in nectar.

As far as we are aware, only one study has investigated the microbiota composition in the crop in *Ae. aegypti* [20]. Gusmão et al. dissected crop tissues from recently eclosed adult females that had not yet fed on sucrose or blood. They found that the crop was dominated by bacteria in the genera *Serratia* and *Bacillus* as well as yeasts in the genera *Pichia* and *Candida*. They also report acidification of the crop and the production of acid by a strain of *Serratia* isolated from this tissue. In *Aedes albopictus*, Guégan et al. [30] compared the microbiota of the crop with that of the midgut in the same individuals and found that alpha and beta diversity of bacterial and fungal communities did not significantly differ between the tissues. They reported that bacteria from the families Weeksellaceae and Burkholderiaceae were most abundant in both crops and midguts in female *Ae. albopictus* and that Corynebacteriaceae was more abundant in crops than in midguts.

In the present work, we used culture-dependent (culturing bacteria on growth media) and culture-independent (qPCR and high-throughput amplicon sequencing of the bacterial 16S rRNA gene) methods to quantify and profile the bacterial communities in paired crops and midguts from adult *Ae. aegypti* females.

## Methods

### Mosquito rearing

*Aedes aegypti* Singapore (Sing) mosquitoes were established from larvae collected in the field in Singapore in 2010 [31]. For strain maintenance, we reared Sing strain larvae at 27°C and 80% residual humidity on a 14:10 light:dark photocycle. We reared larvae in reverse osmosis (RO) water with ad libitum access to larval food (liver powder, tropical fish flake food, and rabbit food pellets mixed in a 2:1:1 ratio and autoclaved) and provided adults with ad libitum access to 10% sucrose. To rear mosquitoes for experiments, we hatched eggs in a vacuum in RO water. When rearing mosquitoes for the culture-dependent experiment, we rinsed eggs one time with 3% bleach, then twice with RO water before hatching. In all cases, we thinned larvae to approximately 250 larvae per pan and provided ad libitum access to larval food. Adults were maintained on 10% sucrose until dissection.

### Dissection and sample preparation

We dissected adult females at 4-6 days post eclosion. Mosquitoes used for culture-independent and those used for culture-dependent experiments came from separate batches of mosquitoes. In both cases, we externally sterilized females with 70% EtOH for one minute, then rinsed twice with filter-sterilized 1X phosphate buffered saline (PBS). We dissected midguts and the ventral diverticula of the foreguts (i.e. crops) from adult mosquito bodies in sterile 1X PBS on glass slides (pre-sterilized with 70% EtOH) and moved the tissues to a sterile pool of 1X PBS. Using sterile forceps, we separated the crop at the diverticular valve. The midgut was separated from the crop immediately anterior to the proventriculus and the proventriculus was included with the midgut. We rinsed both tissues twice in separate pools of sterile 1X PBS. Between each step of the dissection (e.g., after removing the digestive tract, after transferring the crop, after transferring the midgut), we cleaned the forceps with 70% EtOH.

### Culture-dependent experiments

#### Sample processing

We dissected midguts and crops from eight adult females as above and transferred each to separate 1.5ml microcentrifuge tubes containing 150μL 1X PBS without pooling (8 crop samples and 8 paired midgut samples). We kept all samples on ice until homogenizing them using a sterile pestle. We then serially diluted all samples 10^−2^ and 10^−4^ with additional sterile 1X PBS and spread 50μL of each dilution plus the undiluted sample on tryptic soy agar (TSA) and M9 minimal media using sterile beads. Plates were incubated at room temperature for 72 hours and then transferred to 4°C.

#### Bacterial colony characterization and quantification

After 72 hours of bacterial growth, we counted colony forming units (CFUs) and grouped them by colony type using morphological characteristics (e.g., size, color, border, elevation). Counts were made on the lowest dilution plate where CFUs could be readily identified (generally <100 CFU/plate) and back calculated to determine the total number of each colony type present within each tissue. We re-isolated each colony type on fresh media and sequenced the 16S rRNA gene by colony PCR using primers 27F and 1492R [32].

We manually trimmed sequences for each colony type to remove low quality sections and then aligned forward and reverse sequences using the BioEdit Sequence Alignment Editor([33], v7.2.5). If sequences were too short to be aligned or we were only able to obtain high quality sequence in one direction, we used partial sequences. We then used NCBI Nucleotide BLAST [34] with the Megablast algorithm and default parameters to query the Nucleotide Collection (nr/nt) to identify each colony type. We assigned colony type identities using the result with the highest percent identity as long as both the percent identity and query cover were higher than 98%. In two cases, we limited our identification to only the family level (Enterobacteriaceae) because the results showed multiple genera with >98% identity and query cover. All colony types with sequences ≥ 97% similar were combined into a single operational taxonomic unit (OTU), otherwise they were assigned an arbitrary number to indicate different OTUs (e.g., Enterobacteriaceae 1). Abundances and sequences of each colony type can be found in Supplementary File 1.

### Culture-independent experiments

#### Sample processing

For culture-independent experiments, we dissected crops and midguts from eight individuals and pooled each tissue type. This was repeated three times (24 crops and 24 paired midguts, pooled to make 3 total pools of 8). In addition, we dissected crops and midguts from five individuals that were not pooled (5 crop samples and 5 paired midgut samples). We also collected two contamination control buffer blanks that consisted of lysis buffer handled identically to an experimental sample but without tissue added. We placed all samples in 200μL lysis solution from the ZymoBIOMICS DNA Miniprep Kit (Zymo Research, Irvine CA, USA) in 1.5mL microcentrifuge tubes on ice and froze them at −80°C until DNA extraction. We extracted DNA using the ZymoBIOMICS DNA Miniprep Kit according to the manufacturer’s instructions with the following adjustments: all samples were homogenized manually using sterile pestles treated with DNA Away (Thermo Fisher) and rinsed with sterile water, and we eluted DNA in 75μL filter-sterilized water.

#### qPCR

To assess the total amount of bacteria in each tissue sample, we conducted qPCR targeting the 16S rRNA gene as described in [35]. Briefly, each reaction contained 7.5 μL SYBR master mix (Applied Biosystems), 0.35 μL of each primer (primer starting concentrations were all 10 μM), 5 μL template (diluted 1:20), and MilliQ water to a final volume of 15 μL. qPCR conditions were as follows: 95°C for 10 min, (95°C for 15 s then 60°C for 1 min) × 40 cycles. A melt curve was performed after all reactions to verify single product amplification. We estimated 16S copy number of each sample using a standard curve generated from serial dilutions of an *Escherichia coli* 16S PCR amplicon stock of known concentration. To amplify 16S from *E. coli*, we used the same 16S qPCR primers as for the experimental samples. Universal primers used for 16S qPCR are described in [36].

#### High-throughput 16S amplicon sequencing and data processing

High-throughput 16S amplicon sequencing was performed as described in [35] at the Microbial Analysis, Resources, and Services center at the University of Connecticut. Briefly, 5ng of DNA was used as template in a PCR with 515F and 806R primers with Illumina adapters and eight basepair dual indices [37,38]. Many samples (Supplementary File 1) did not amplify using the standard 30 cycles, and in these cases, an additional five cycles were performed which resulted in successful amplification. The contamination control buffer blanks failed to amplify but were still included in the sequencing reaction to account for any potential contamination. 250bp paired-end sequencing was performed on a MiSeq system using the MiSeq Reagent Kit V2 (Illumina, Inc.).

Sequences were demultiplexed using onboard bcl2fastq. We completed all bioinformatic analysis using R software v4.1 [39]. We used the *DADA2* R package v1.16 [40] with default parameters specified by the author ([41]; https://benjjneb.github.io/dada2/tutorial.html) to identify amplicon sequence variants (ASVs), assign each ASV a taxonomic identity using the RDP classifier with training set v18 [42], and generate count data for each ASV. The specific code we used for sequence processing, downstream bioinformatics, and statistical analyses can be found in Supplementary File 2. We imported the data to the *phyloseq* R package v1.30.0 [43] for downstream bacterial community analyses and visualization. We created a phyloseq object from the ASVs identified by DADA2. We then removed ASVs not from the Bacteria kingdom and those classified as either mitochondria or chloroplast. As low biomass samples are prone to the effect of environmental and reagent contamination [44,45], two buffer blank controls were included in the experimental design. Before any other filtering step was applied to the phyloseq object, we explored the composition of these controls and their similarities to the experimental samples to better understand the nature of potential contamination and address it. The most abundant taxon in the buffer blank controls was *Acinetobacter*, and *Elizabethkingia* was present as well. These taxa are associated with *Aedes* mosquitoes [46] and their natural environment [10,47,48] but are also known laboratory contaminants and are commonly found in this environment [45,49]. The average number of reads in the controls accounted for a very low percentage (0.51%) of the mean reads per sample in our dataset (Supplementary Figure 1), and a principal coordinate analysis (PCoA) revealed that buffer blank controls clustered separately from experimental samples (Supplementary Figure 2). Therefore, we proceeded with the analyses despite this minor contamination. As an additional quality control step, we used the *decontam* R package v1.6 [50] with default prevalence filtering parameters to remove ASVs identified as contamination. Buffer blank control samples were then removed from the dataset. Finally, we removed ASVs accounting for fewer than 5 total reads and then scaled reads per sample based on the smallest read count as suggested by [51] in order to preserve proportions.

### Statistical analysis

For both the culture-dependent experiments and the qPCR of the culture-independent experiments, we assessed the differences in microbiota size between the crop and midgut tissues. We used either a t-test or Kruskal-Wallis test in R (v4.1, [39]) depending on whether the data conformed to the assumptions of a parametric test. We also assessed whether the variance of the microbiota in each tissue was significantly different using Levene’s test in the *car* package v3.0-13 [52].

#### Microbial ecology analyses

Unless stated, we used R (v3.3.2) and the Rstudio environment [53] to perform all the statistical analysis and graphical representations for the high-throughput amplicon sequencing analysis. To measure alpha diversity, we calculated the observed richness and the Gini-Simpson index (1-D) [54] for each sample with *phyloseq*. Differences between tissues were evaluated by a Kruskal-Wallis test separately for each sample type (single and pooled) using *ggpubr* v0.4.0 [55]. We assessed beta diversity and clustering profiles using non-constrained (PCoA) and constrained ordinations (Canonical Correspondence Analysis, CCA) based on Bray-Curtis dissimilarities. The PCoA plot was generated with the *microeco* package v0.3.3 [56] and the CCA plot with the *vegan* package v2.5.7 [57]. To partition the variance within the data matrix, we used the adonis2 function in *vegan* to run a permutational multivariate analysis of variance (PERMANOVA) to test the effects of tissue (crop or midgut), sample type (single or pooled), and the most abundant taxa in the dataset on variation in bacterial community composition. We then applied two approaches to identify microbial community members significantly associated with each tissue type. We chose a combination approach as recommended by [58] to prevent misidentifying discriminant features due to technical and analytical artifacts (e.g. sequencing depth and filtering of rare ASVs). The first, LEfSE [linear discriminatory analysis (LDA) effect size] [59], identifies biomarkers that are differentially abundant between treatments (tissue type in our case) and ranks them based on their association with each treatment level. The analysis was run with an LDA cutoff value of 3.0 and p-value of 0.05 using the *microeco* package. The second method, indicator species analysis, relies on community ecology principles to identify taxa reflective of their niche state and predicts the diversity of other community members. We searched for indicator ASVs using the multipatt function within the *indicspecies* package v1.7.0 [60]. To perform functional profile predictions of the prokaryotic communities within each tissue, we used FAPROTAX [61]. This algorithm maps the ASV compositional profile of each sample against a curated database of metabolic and ecologically relevant functions based on literature associated with cultured bacterial representatives. We tested the significance of the functional profile differences between tissue groups with a Wilcoxon rank-sum test using the *ggpubr* package reporting p values < 0.05 after a false discovery rate correction.

## Results

### Size of the microbiota is more variable in midguts compared to crops

We first estimated the number of bacteria in each sample by quantifying 16S rRNA gene copy number using qPCR. Pooled samples showed a different pattern than single (non-pooled) samples. The variance was highly similar between tissues for pooled samples (Levene’s test for equal variances: F = 0.256, p = 0.640) but in single samples, the variance of midguts was significantly larger than crops (Levene’s test for equal variances: F = 6.12; p = 0.038). Pooled midgut samples trended toward a higher log_10_ 16S copy number (median_midgut_p_ = 5.05) than pooled crop samples (median_crop_p_ = 3.98), though the tissues were not significantly different using a paired t-test (t = 3.69, p = 0.066, Figure 1). The median copy number for single midgut samples was highly similar to that of single crop samples (median_midgut_s_ = 2.48; median_crop_s_ = 2.79), and tissues were not significantly different using a Wilcoxon rank sum test (p = 0.691, Figure 1).

**Figure 1:**
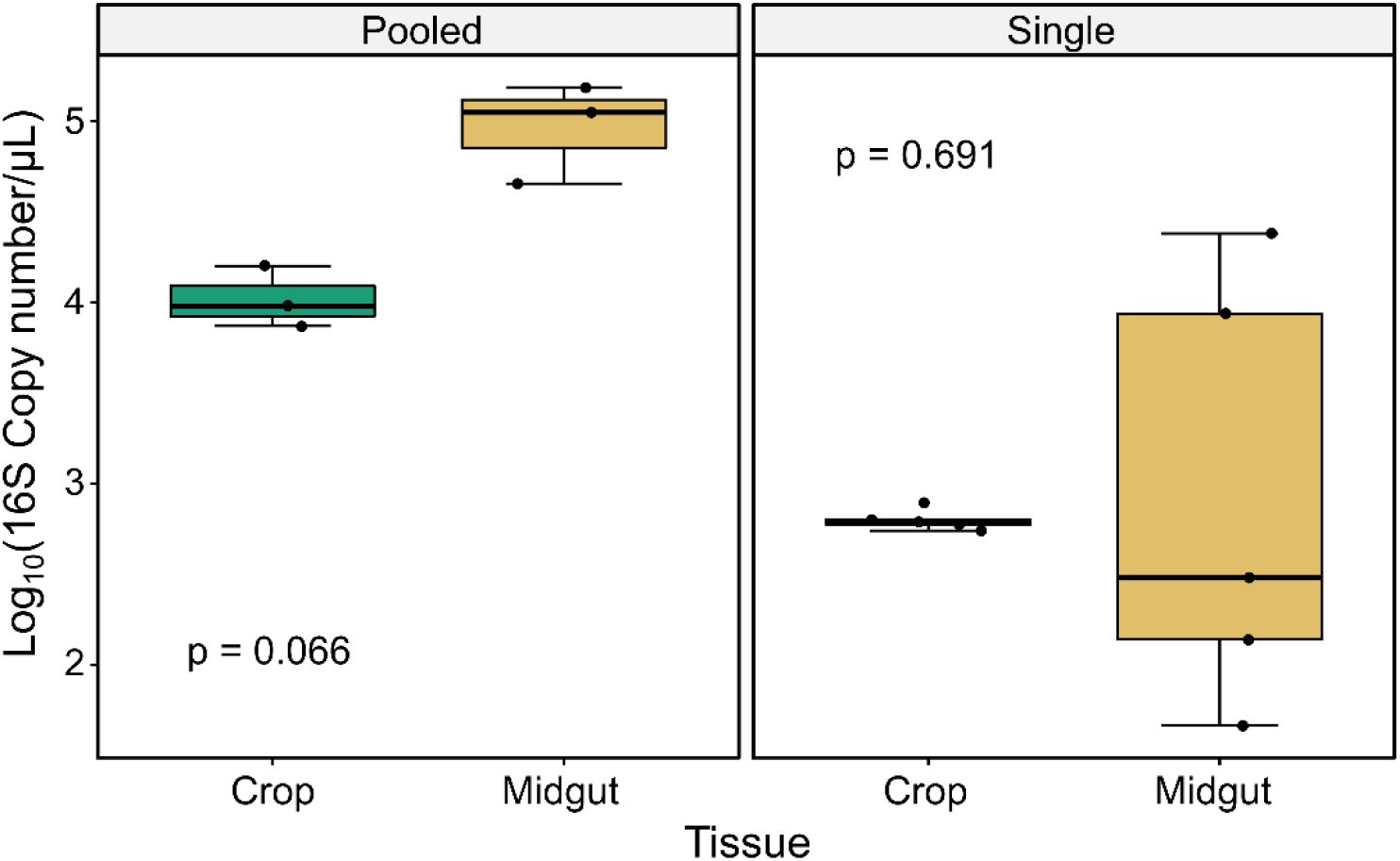
Microbiota size measured using qPCR is more variable in midguts compared to crops. We measured copy number of the 16S bacterial rRNA gene using qPCR. Samples were either pooled (left panel) in groups of 8 (n = 3 pools per tissue type) or assayed as individual single tissues (right panel) (n = 5 samples per tissue type). For pooled samples, variance was similar between tissues (Levene’s test for equal variances: F = 0.256, p = 0.640), but for single samples the variance among midgut samples was significantly larger than crop samples (Levene’s test for equal variance, F = 6.12, p = 0.038). For pooled samples, median log_10_(16S copy number/μl) trended higher in midguts (median = 5.05) than in crop samples (median = 3.98) though the difference between tissues was not significant using a paired t-test (t = 3.69, p = 0.066). For single samples, median values were similar for midguts (median = 2.48) and crops (median = 2.79) and were not significantly different using a Wilcoxon rank sum test (p = 0.691).

We also estimated the total number of cultivable bacteria in each tissue by homogenizing single, paired crops and midguts from adult female *Ae. aegypti* and culturing the homogenate on TSA and M9 media at multiple dilutions. Variances were significantly different between tissues for both media types as measured by Levene’s test (TSA: F = 5.521, p = 0.034; M9: F = 11.554, p = 0.004). The median number of CFU/sample trended higher in midguts (median_midgut_TSA_ = 4.0×10^6^, median_midgut_M9_ = 6.5×10^6^) than in crops (median_crop_TSA_ = 1.9×10^3^, median_crop_M9_ = 2.9×10^3^, Figure 2) for both media types, but these differences were not statistically significant as measured by a Kruskal-Wallis test (p_TSA_ = 0.074, p_M9_ = 0.093).

**Figure 2:**
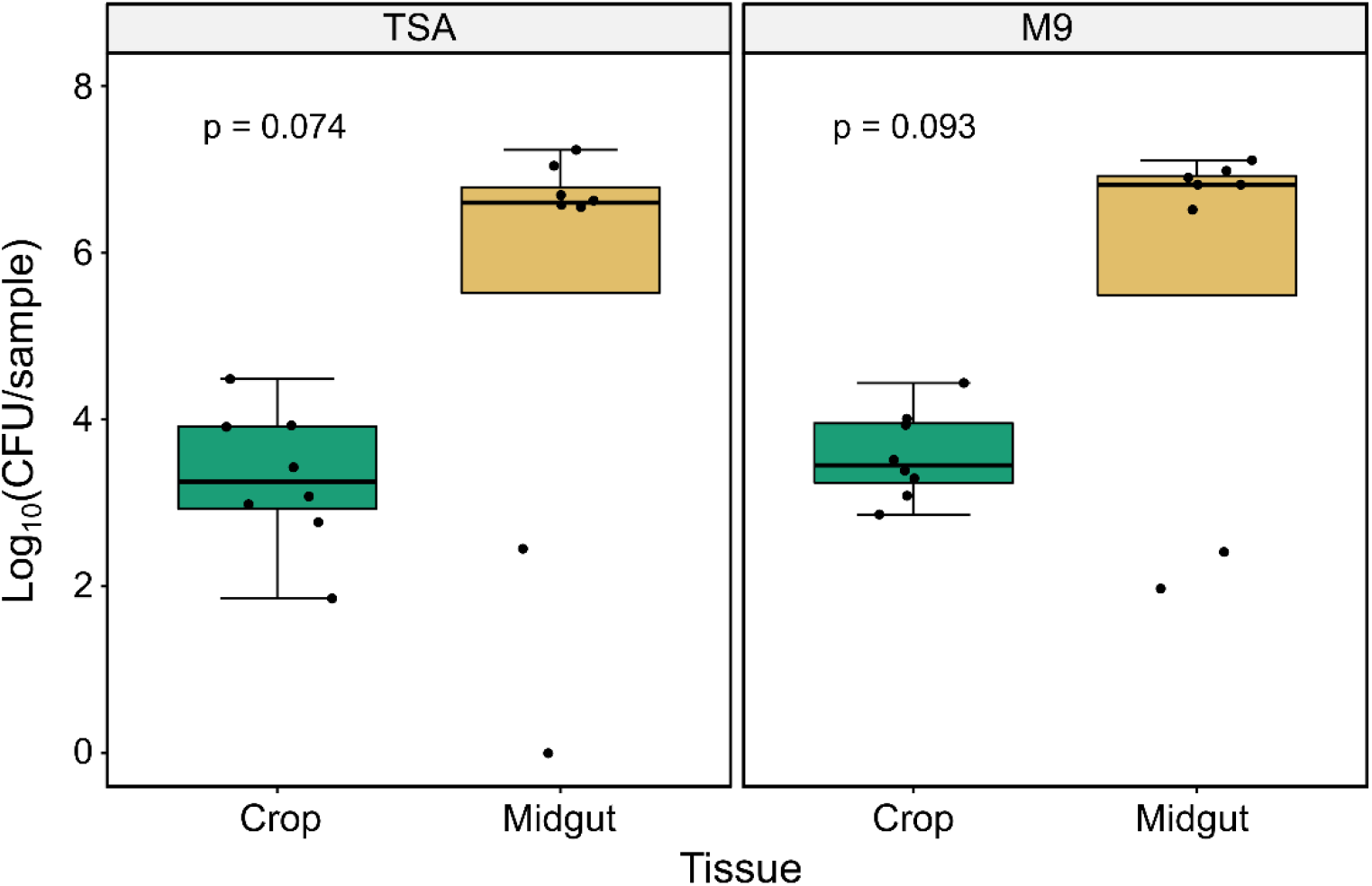
Bacterial load in crops and midguts of adult female *Ae. aegypti*. Individual crops and midguts from eight adult females were homogenized and cultured on TSA (left panel) and M9 (right panel) media. Variance was significantly different between tissues for both media types (TSA: F = 5.521, p = 0.034; M9: F = 11.554, p = 0.004) as measured by Levene’s test for equal variances. Tissue type was not a significant predictor of bacterial load on either media (TSA: p = 0.074; M9: p = 0.093) as measured by Kruskal-Wallis test.

### Community composition of crop and midgut tissues using culture-independent and culture-dependent approaches

We sequenced the bacterial community of 16 samples (3 pools of 8 crops, 3 pools of 8 midguts, 5 individual crops, and 5 individual midguts) and 2 buffer blank controls resulting in 534462 amplicon reads. One individual midgut sample failed during sequencing and no data were obtained for that sample. After quality filtering, decontamination, and scaling to address the effect of sequencing depth, we retained 9221 reads as the minimum read number per sample. In total, the dataset comprised 134 ASVs taxonomically distributed into 10 bacterial phyla. From them, 97% were classified within 6 phyla as follows: Proteobacteria (48.8%), Firmicutes (23.7%), Actinobacteria (11.1%), Chloroflexi (6.6%), Planctomycetes (3.7%), and Bacteroidetes (2.9%).

All crop samples (pooled and single) were dominated by bacteria from the family Acetobacteraceae. Bacteria from the family Weeksellaceae were also present in appreciable amounts in many crop samples but never reached higher than 9.8% of the total community (Figure 3). Midgut samples were dominated by Weeksellaceae when pooled, but single samples were more variable. Two of four single midguts had near total dominance of Weeksellaceae while the other two midguts contained Acetobacteraceae and multiple other families (Figure 3).

**Figure 3:**
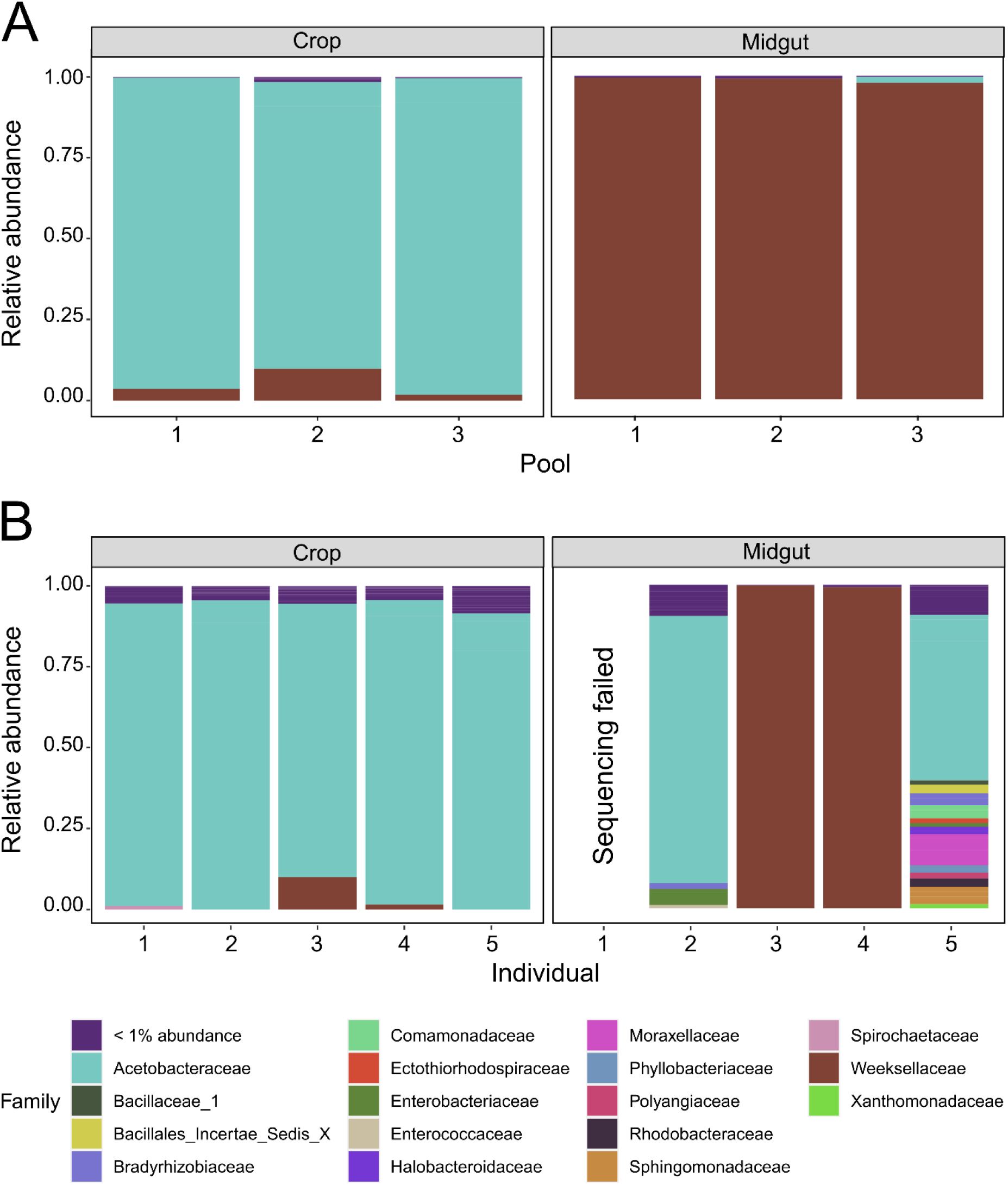
Composition of microbial communities in crops and midguts of *Ae. aegypti* adult females using a culture-independent approach. Relative abundances of bacterial families are clustered by tissue (labeled across the top) and sample type. In (A), samples are pooled in three groups of eight. Tissues from each numbered pool are paired, i.e., they were collected from the same eight individuals. In (B), samples are single (un-pooled) paired tissues, i.e. each numbered crop came from the same individual as the matching numbered midgut. Bacterial communities in crops are dominated by Acetobacteraceae. Midguts are mostly dominated by Weeksellaceae, though individual midgut samples 2 and 5 lack Weeksellaceae and are dominated by Acetobacteraceae.

We also evaluated community composition by culturing 8 single, paired crops and midguts on TSA and M9 media (Figure 4). These were different individuals than those used in Figure 3. We characterized and counted bacterial CFUs based on morphological characters, then sequenced the 16S rRNA gene from a representative isolate to identify each colony type to genus. Samples cultured on both types of media showed similar, but not identical, results. The largest difference between media types was that we did not observe *Tanticharoenia* bacteria on TSA media (Figure 4A) but did using M9 media (Figure 4B). Similar to what was observed in the culture-independent experiments, genera in the family Acetobacteraceae (i.e., *Asaia* and *Tanticharoenia*) dominated the crop samples (Figure 4A, B). *Elizabethkingia* (Family Weeksellaceae) were present in 75% of midgut samples and represented more than 99.9% of observed CFUs when present (Figure 4A, B). *Elizabethkingia* were also dominant in 37.5% of crops and present in varying amounts in all but one. However, *Elizabethkingia* made up a smaller proportion of CFUs in crops compared to midguts (Figure 4A, B).

**Figure 4:**
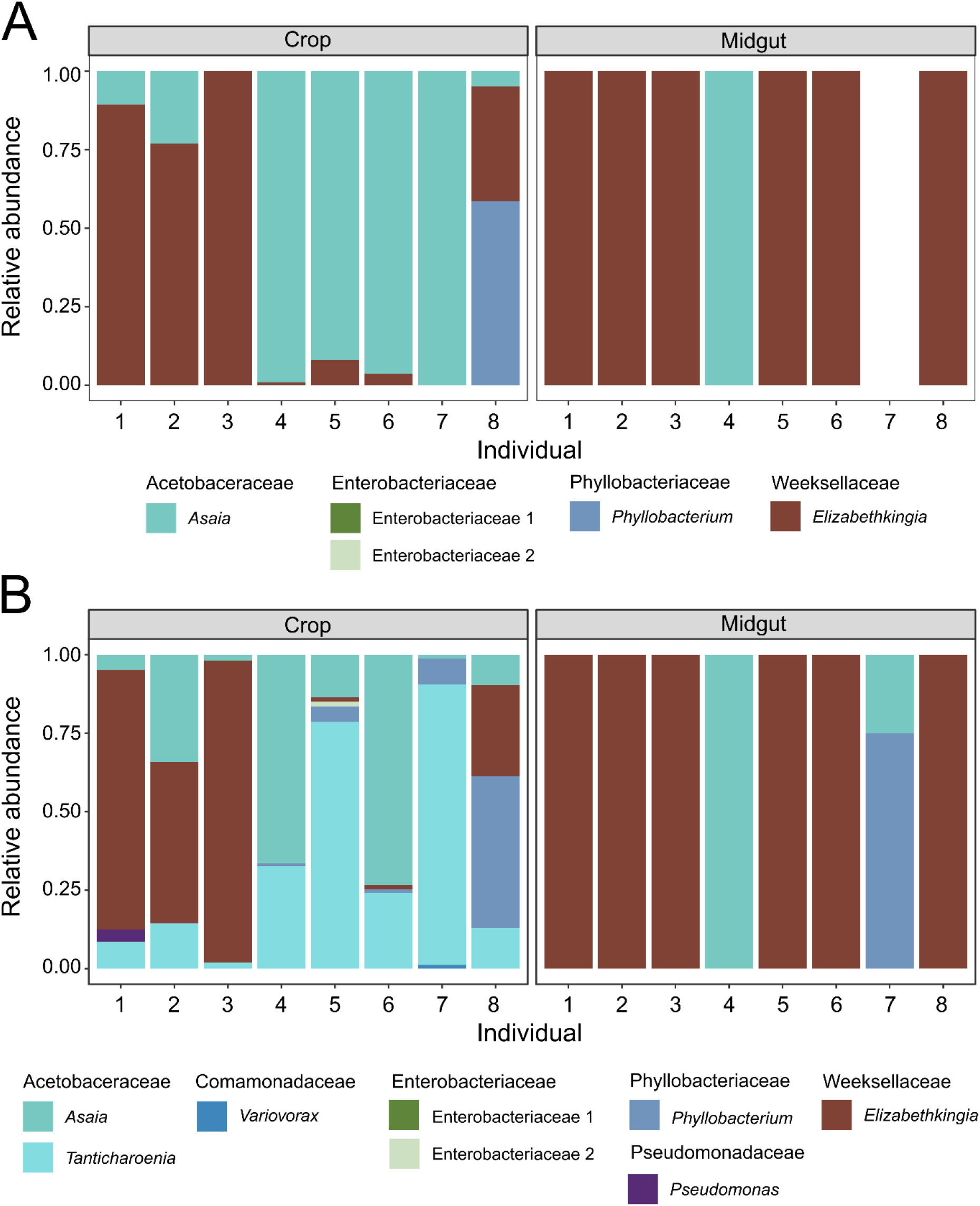
Composition of microbial communities in crops and midguts of *Ae. aegypti* adult females using a culture-dependent approach. Relative abundances of bacterial genera are clustered by tissue (labeled across the top). Individual crops and midguts from eight adult females were homogenized and an aliquot from each was then cultured on TSA (A) and M9 (B) media. In all cases, samples are single (un-pooled) paired tissues, i.e., each numbered crop came from the same individual as the matching numbered midgut. Bacterial communities in crops are dominated by *Asaia, Tanticharoenia*, and *Elizabethkingia*. Midguts are almost all dominated by *Elizabethkingia*, though individual midgut samples 4 and 7 lack this genus.

### Bacterial community diversity analysis in culture-independent experiments

Following our initial observations of microbial community composition, we next analyzed the high-throughput amplicon sequencing data to evaluate whether alpha and beta diversity differ by tissue type and whether any specific bacterial taxa are significantly associated with crops and/or midguts.

#### Bacterial communities within pooled crop samples are more diverse than midguts

For pooled samples, alpha diversity measured as observed ASV number (Figure 5A, p = 0.05) and Gini-Simpson index (Figure 5B, p = 0.05) was marginally higher in crops than midguts when evaluated by a Kruskal-Wallis test. Using Levene’s test for equal variances, we determined that the variance of pooled samples was similar between tissues for observed ASV number (F = 2.28, p = 0.205) but higher in crops than midguts for Gini-Simpson (F = 10.08, p = 0.03). For single samples, alpha diversity was not significantly different between tissues, but Levene’s test showed much higher variance among midgut samples than crops for observed ASV number (F = 18.70, p = 0.003) and Gini-Simpson (F = 9.92, p = 0.001).

**Figure 5:**
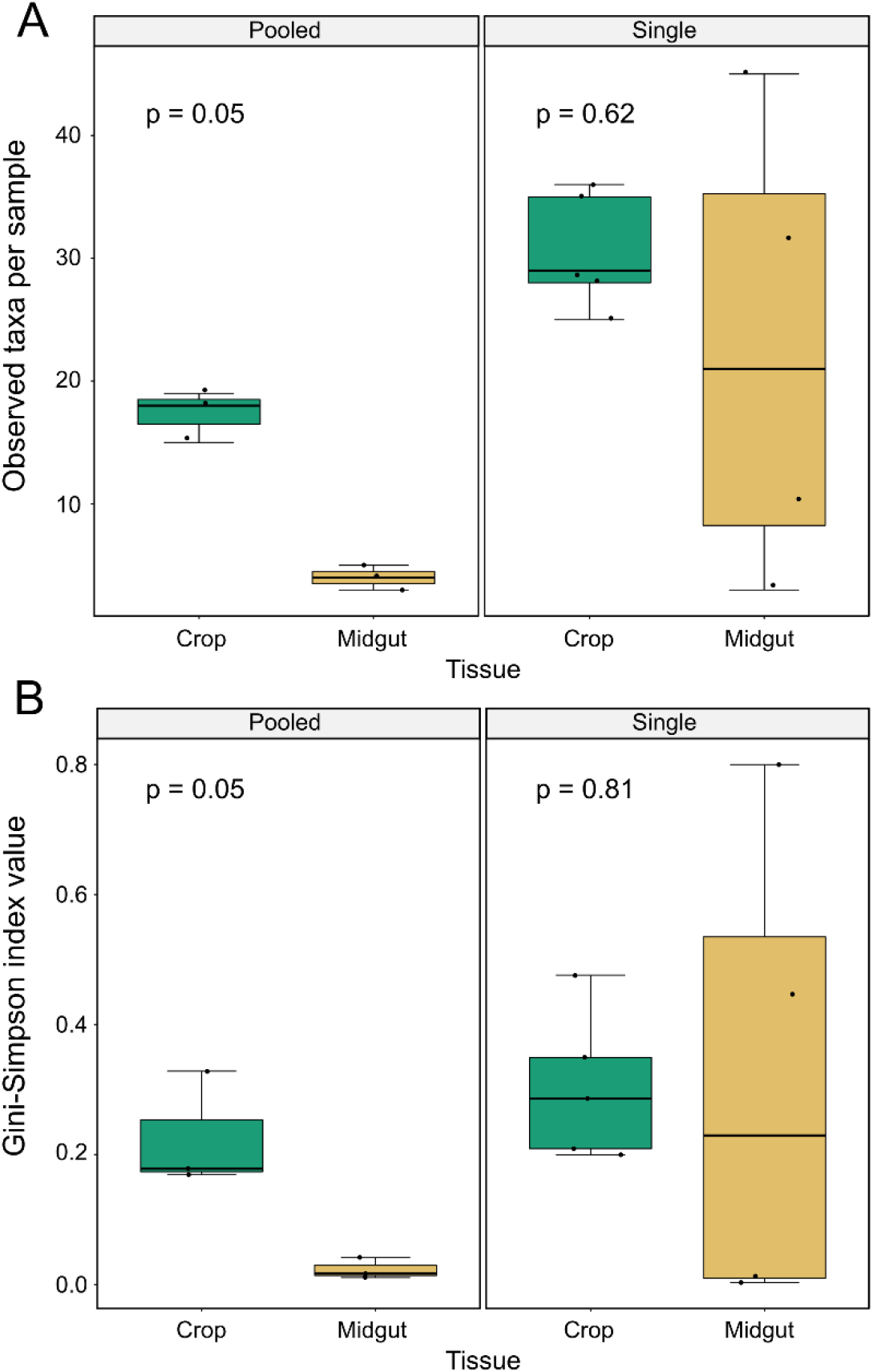
Alpha diversity differs between crop and midgut samples. We measured alpha diversity as observed ASVs (A), and the Gini-Simpson index (B). For pooled samples (left panels), both measures of alpha diversity were marginally significantly higher in crops than midguts (Kruskal-Wallis test, p = 0.05 for both measures) Variance among crops was significantly larger than midguts for Gini-Simpson indices (Levene’s test: F = 10.08, p = 0.03) but not for observed taxa values (Levene’s test: F = 2.28, p = 0.205). For single samples (right panels), tissues were not significantly different for either measure, but midgut samples showed a much larger range of values than crops for both observed taxa values (Levene’s test: F = 18.70, p = 0.003) and Gini-Simpson indices (Levene’s test: F = 9.92, p = 0.001).

#### Composition and structure of microbiota in crops significantly differs from that in midguts

We executed a PCoA on the high-throughput amplicon sequencing data to explore how tissue (crop and midgut) and sample type (pooled or single) drove the community profile variation among samples (Figure 6A). The first two components encompass 98.2% of the total variance (91.7% PCoA1 and 6.5% PCoA2). Samples predominantly grouped in homogeneous clusters by tissue type, separating themselves along axis 1. Two midgut samples digressed from this diversity profile and clustered separately. These two data points represent single midguts samples 2 and 5 (Figure 3) which had low numbers of Weeksellaceae and a larger number of Acetobacteraceae. In order to test the significance of tissue and sample type as driving variables of beta diversity, we performed a PERMANOVA. This analysis revealed tissue as the only significant variable (R^2^ =54.81 p = 0.003). Sample type accounted for a non-significant 6.6% of variance (p = 0.107), and no significant interaction among the effector variables was detected. The communities were dominated by reads that map to the genera *Elizabethkingia* (Family: Weeksellaceae), *Tanticharoenia* (Family: Acetobacteraceae), and/or *Asaia* (Family: Acetobacteraceae) (Figure 3). Therefore, we conducted a CCA (Figure 6B) to address how the three most abundant ASVs in the dataset (which map to *Tanticharoenia, Elizabethkingia*, and *Asaia*) may be driving the diversity profiles detected. *Elizabethkingia* reached maximum abundance in midgut samples, whereas both *Tanticharoenia* and *Asaia* were maximally abundant in crops. Tight clustering of samples suggests that tissue and dominant taxa included in the analysis account for a large amount of variance. A second PERMANOVA considering abundance counts of *Elizabethkingia*, *Tanticharoenia*, and *Asaia*, and their interactions with tissue type as variables (Table 1) confirmed that the variance is significantly driven by these dominant taxa and their respective interactions with tissue type, accounting for 99.68% of the total beta diversity.

**Figure 6:**
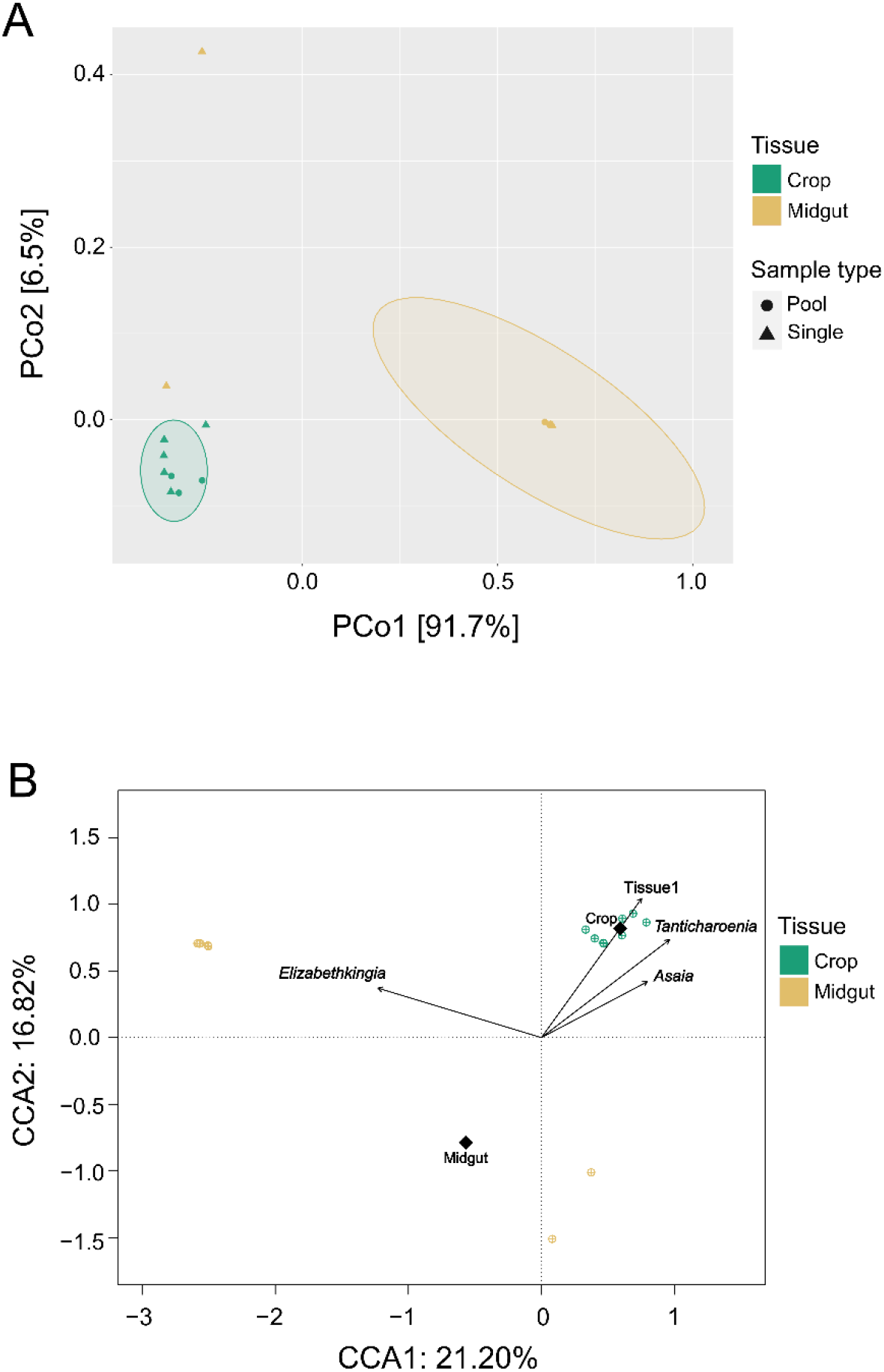
Ordination analyses reveal significantly different bacterial community structures between crop and midgut tissues. Beta diversity was estimated using Bray-Curtis dissimilarities. The unconstrained analysis (principal coordinates analysis, PCoA) (A) shows the samples cluster by tissue type predominately along axis 1 (91.7%). The constrained analysis (canonical correspondence analysis, CCA) (B) shows that the explanatory variables (Table 1) capture 21.2% and 16.82% of variation in CCA1 and CCA2, respectively. The group centroids of each tissue (represented by a black diamond) are located in opposing coordinates, indicating a difference between the communities. The abundance of the dominant taxa (*Asaia, Tanticharoenia*, and *Elizabethkingia*), shown as vector arrows, indicate that *Asaia* and *Tanticharoenia* are maximally abundant in crops while *Elizabethkingia* is maximally abundant in midguts.

**Table 1:** A PERMANOVA to test for the association between beta diversity (quantified by Bray-Curtis dissimilarities) and explanatory variables representing tissue type and the raw abundance of the three dominant taxa in the data set.

| Explanatory variable | R<sup>2</sup> | p – value |
| --- | --- | --- |
| Tissue | 0.0291 | 0.001 |
| *Asaia* | 0.0281 | 0.001 |
| *Elizabethkingia* | 0.7446 | 0.001 |
| *Tanticharoenia* | 0.0439 | 0.001 |
| Tissue:*Asaia* | 0.0959 | 0.001 |
| Tissue:*Elizabethkingia* | 0.0048 | 0.001 |
| Tissue:*Tanticharoenia* | 0.0501 | 0.001 |
| Residuals | 0.0104 |  |
| Total | 1.00000 |  |

#### Elizabethkingia and Tanticharoenia are biomarkers of midgut and crop tissues, respectively

We used two methods to test for differentially abundant taxa with significant associations to each tissue. Linear discriminant effect size values estimated with the LEfSe tool (Figure 7) detected ASVs within the Bacteroidetes clade as consistently more abundant in the midgut. This was driven by ASV_2 and ASV_281, which were classified as *g_Elizabethkingia* (f_Weeksellaceae) and an unidentified member of f_Weeksellaceae, respectively, (LDA = 3.48, p < 0.01). LEfSe also detected ASVs within the Proteobacteria clade as biomarkers for the crop. Within this clade, ASV_1 (the most abundant amplicon in the dataset), was classified as *g_Tanticharoenia* (f_Acetobacteraceae), had an LDA of 3.49 (p < 0.01). Although below the LDA score cutoff, ASV_3, identified as *g_Asaia* (f_Acetobacteraceae) was also detected as a significant crop biomarker (LDA = 2.33, p < 0.01). Indicator species analysis detected two ASVs as reliable predictors for the crop. One was ASV_1 (g_*Tanticharoenia*) and the second was a low-abundance ASV that mapped to *g_Rubrobacter* (Table 2). No significant indicator ASVs were identified for the midgut using this method (Table 2).

**Figure 7:**
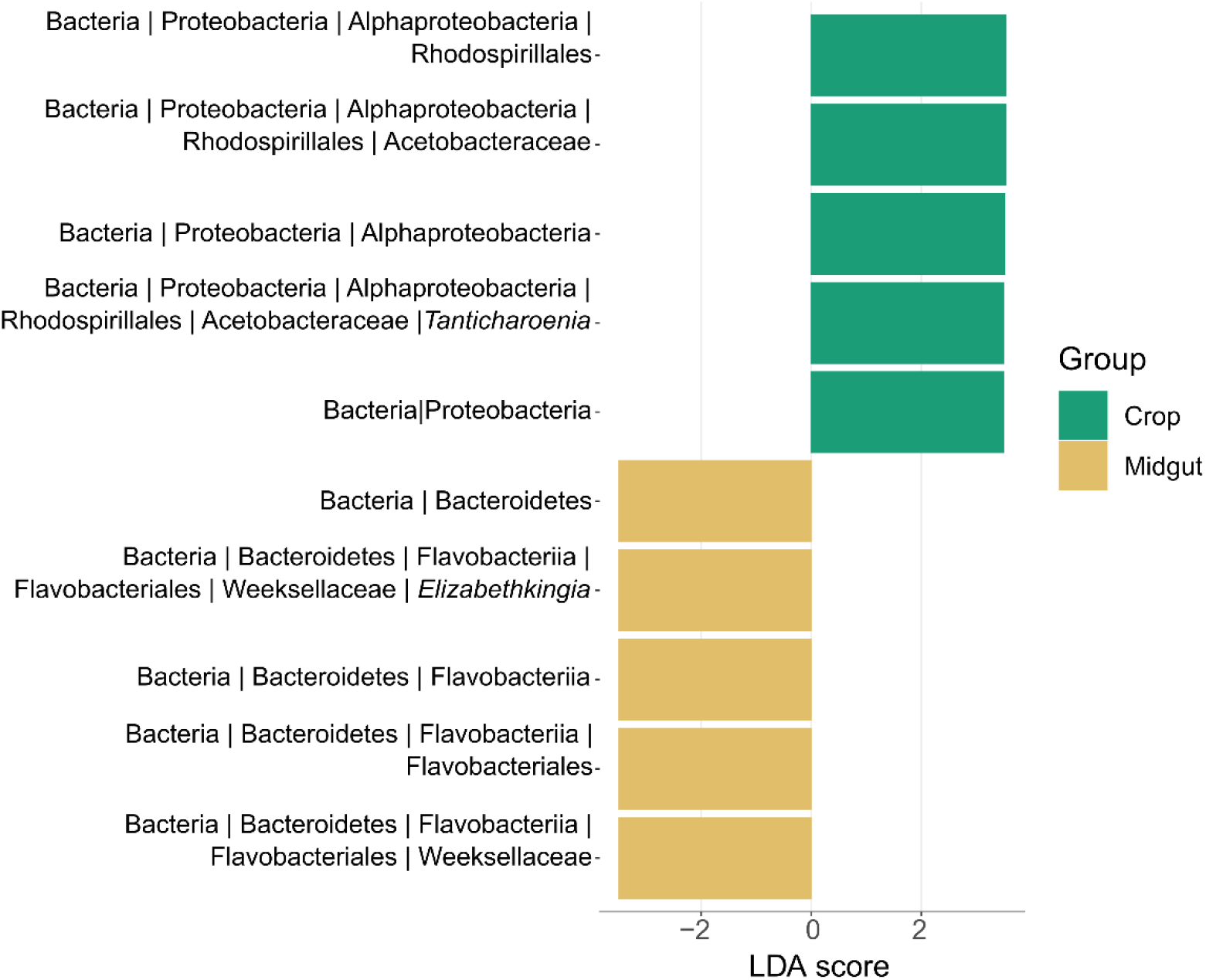
Tissue specific biomarkers determined by LDA scores using LEfSe method. At an LDA cutoff point of 3 (p < 0.01), two bacterial clades representing three ASVs were identified as significantly differentially abundant between tissues.

**Table 2.**
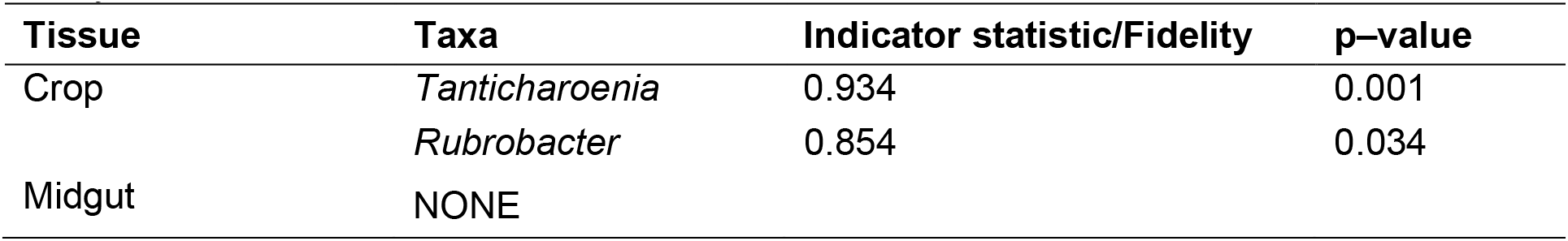
ASVs significantly associated with tissue samples as revealed by indicator species analysis.

| Tissue | Taxa | Indicator statistic/Fidelity | p-value |
| --- | --- | --- | --- |
| Crop | *Tanticharoenia* | 0.934 | 0.001 |
|  | *Rubrobacter* | 0.854 | 0.034 |
| Midgut | NONE |  |  |

#### Functional prediction analysis reveals potential differences in microbial community metabolic function between crop and midgut

FAPROTAX prospection, a tool used for metabolic phenotype predictions, revealed 34 functional categories represented in the microbial communities in the full dataset (Figure 8). Among them, 15 were shared between both tissues. Fourteen functional categories were only found in crops, including methanotrophy, methanol oxidation, methylotrophy, chitinolysis, knallgas bacteria, dark hydrogen oxidation, and hydrocarbon degradation, and five functional categories were only found in midguts, including manganese oxidation. Of the 19 categories found in only one tissue, there were 11 singletons (inferred from only one sample). Chemoheterotrophy and aerobic chemoheterotrophy were ubiquitous among all samples (Figure 8). Chemoheterotrophy was the most abundant functional category in both tissues, accounting for 64.38 ± 3.00% in the crop and 48.96 ± 5.00 % in the midgut. Comparative analysis revealed two functional categories that were significantly more represented in crops than midguts: methylotrophy, which is binned within the carbon metabolism related functions in the FAPROTAX database, and methanol oxidation, which is binned in the broader “other annotated categories” in the database (Figure 8). Though not significant, fermentation, aromatic compound degradation, nitrate reduction, and animal parasites/symbionts were more commonly observed in crop samples, and at higher community proportions than in midguts (Figure 8).

**Figure 8:**
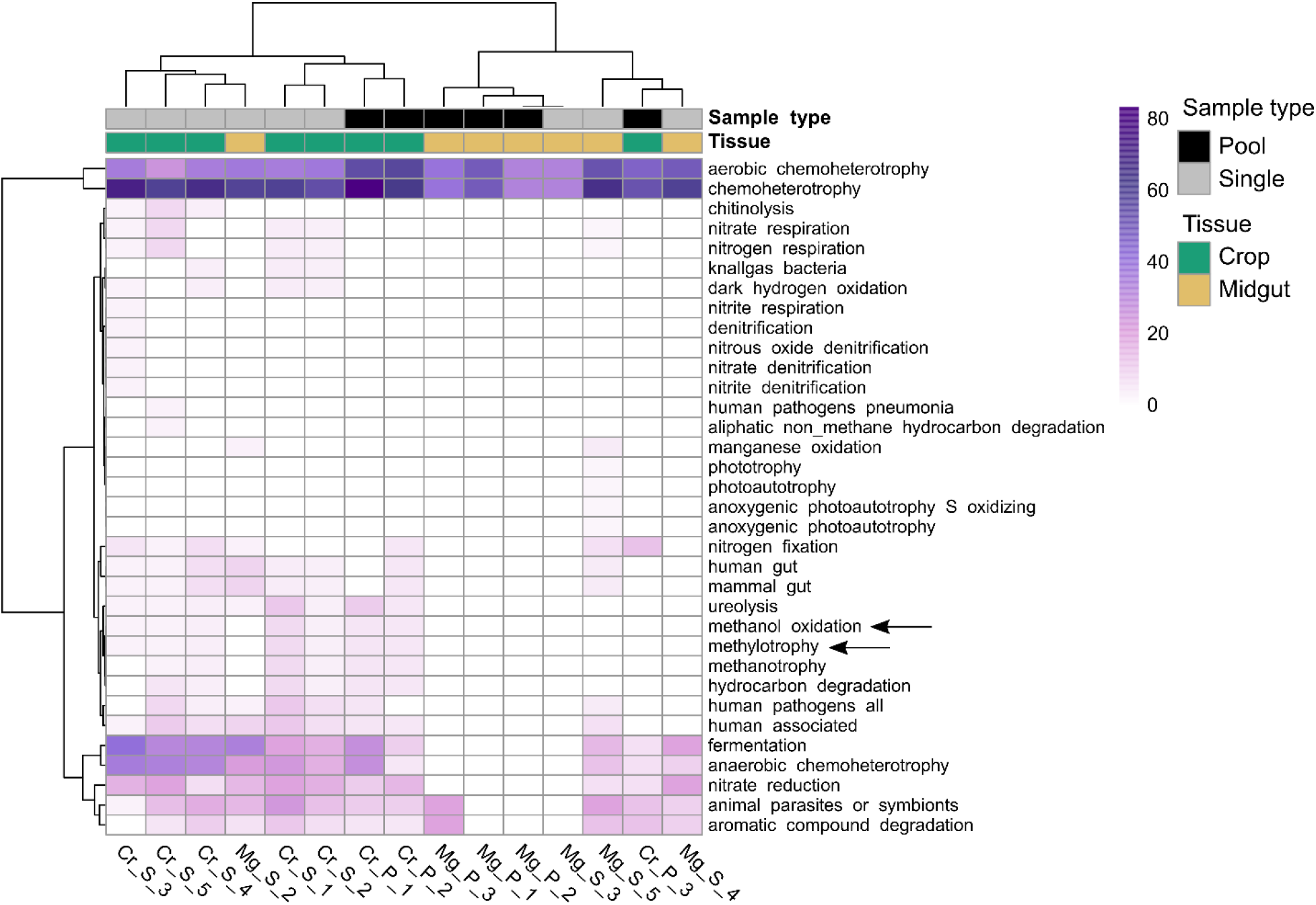
Predicted community metabolic profiles reveal a predominant clustering pattern driven by tissue. The heatmap represents the relative abundance of the predicted pathways (rows) in the community from each sample (columns) based on the FAPROTAX database. The color key highlights the tissue of origin (teal/beige) and sample type (black/gray) of each bacterial community. Statistical significance indicated by an arrow (p < 0.05) was tested by Wilcoxon rank sum test followed by an FDR correction.

## Discussion

In the current work, we used culture-dependent and culture-independent techniques to investigate the bacterial communities found in the crop and midgut of laboratory reared adult female *Ae. aegypti* mosquitoes. Our results reveal similarities and differences between the microbial communities of these two tissues.

The average size of the microbiota was not significantly different between tissues, though we did observe a trend toward crops having smaller and more diverse bacterial communities than midguts. However, microbiota size in midguts was dependent on whether the sample was pooled or single and by the composition of the microbiota. When measured singly, the size of the microbiota in midgut samples was revealed to be highly variable, whereas crop microbiota size was highly consistent between replicate samples in all cases. Moreover, midgut samples with the largest microbial communities were, in all cases, dominated by *Elizabethkingia* (Family: Weeksellaceae), while those with smaller microbial communities lacked *Elizabethkingia*, suggesting the presence or absence of this bacterial genus is a primary determinant of microbiota size in midguts. In contrast, crop samples had highly similar microbial community sizes regardless of composition.

We found that communities in both tissues were relatively simple, consisting of fewer than 20 ASVs. Our richness and within-sample diversity profiles are consistent with the low diversity reported in other studies that have sequenced the midgut and/or crop microbiota of laboratory reared sugar fed *Aedes* mosquitoes (e.g. [6,30,62–65]). Diversity within samples (alpha diversity) was higher in crops than midguts, but again, this was only for pooled samples; when sampled singly, we found that alpha diversity was more variable among midgut samples than among crops, and this was largely driven by the overdominance of *Elizabethkingia* in midguts. Microbial communities differed significantly in composition and structure between tissues (beta diversity). Crop tissues were primarily dominated by bacteria from the family Acetobacteraceae, while midguts were primarily dominated by bacteria from the family Weeksellaceae. The family Weeksellaceae was present in crops but at lower relative abundance than in midguts. Midguts also contained Acetobacteraceae but only in a minority of samples and only when Weeksellaceae was absent. A PCoA confirmed clustering of samples by tissue. A CCA coupled with a PERMANOVA showed that Acetobacteraceae and Weeksellaceae were significant drivers of differences between tissues and that Acetobacteraceae was significantly associated with crop tissues while Weeksellaceae was significantly associated with midguts. A biomarker analysis was consistent with these findings and identified *Tanticharoenia* and *Elizabethkingia* as markers of crops and midguts, respectively. We note that the differential abundance analysis methods differ in sensitivity and specificity, and LEfSE, which is a common method among vector biology microbiome studies, is susceptible to rare feature counts [66] and can result in false positives. Nearing et al. (58) stresses that some methods are reported interchangeably but can considerably differ in power and susceptibility to experimental design. We therefore complemented our LEfSE analysis with an indicator species analysis, which did not give identical results but did corroborate the finding of *Tanticharoenia* as a biomarker for the crop.

Bacteria in the family Acetobacteraceae, specifically *Tanticharoenia* and *Asaia* in our samples, are considered “acetic acid bacteria” (AAB, [67,68]). *Tanticharoenia* was previously denominated as *Asaia* but was recently determined to be a distinct genus [68]. This group of bacteria is characterized by the common ability to catabolize sugars and produce acidic compounds from them (e.g., acetic acid). Specifically, *Tanticharoenia* produces acid from ethanol and multiple sugars, and can grow in the presence of 0.35% acetic acid [69]. Interestingly, *Asaia* is the only Acetobacteraceae that does not produce acetic acid from ethanol but can oxidize acetate and lactate to carbon dioxide [70].

Our midgut samples were highly dominated by *Elizabethkingia* (Family: Weeksellaceae). *Elizabethkingia*, and other bacteria from the phylum Bacteriodetes, are commonly reported in mosquito midguts, especially in laboratory settings but also in wild-caught adult specimens and *Ae. aegypti* larval development sites (e.g. [48,62,71–74]). *Elizabethkingia* has been previously shown to outcompete other bacteria in the midgut [75] and has been observed in midgut tissue of *Ae. aegypti* Singapore strain in high numbers [76]. Therefore, it was not unexpected that this genus was prevalent in our midgut samples; however, we consider it notable that *Elizabethkingia* was less prevalent in crops than in midguts.

Gusmão et al. [20] cultured bacteria from laboratory reared *Ae. aegypti* adult female crops on Brain Heart Infusion agar. They primarily observed yeast species and *Serratia* in *Ae. aegypti* crops. We did not identify *Serratia* in our crop or midgut communities, though we did observe the presence of other Enterobacterales (*Klebsiella* and *Enterobacter*). Additionally, ASV_22 from our study was assigned to Enterobacteriaceae but was not identified at the genus level. Thus, we cannot rule out that *Serratia* is present in our community data set as an unidentified ASV. We also note differences between our findings and those of Guégan et al. [30] who reported no significant differences in alpha diversity and bacterial community structures when comparing crop and midgut samples from *Ae. albopictus*. They found that Weeksellaceae was the most abundant taxa in female crops, and Burkholderiaceae was most dominant in midguts. They did not find any Acetobacteraceae present in any tissue. Differences between their findings and ours may be due to several experimental design differences such as the mosquito species, age, and/or diet of the females. Similarly to our study, the two previous studies of *Aedes* crop microbiota [20,30] were performed in the laboratory, and since mosquitoes primarily acquire their microbiota from the environment [73,77], laboratory rearing inherently influences and limits the potential members of the microbiota to those present in the artificial laboratory environment. This may explain the lack of Acetobacteraceae in crops of *Ae. albopictus*. It would be valuable to assay the crop and midgut microbiotas of field-collected mosquitoes to determine if Acetobacteraceae is common in the crops of wild mosquitoes. Wild mosquitoes will be exposed to a different suite of bacteria in their natural larval development sites and will feed on nectar as adults instead of pure sucrose which is not only nutritionally different but may also contain antimicrobial compounds that can influence the crop bacterial community [78].

One possible driver of the differences we observed in microbial communities between the crop and midgut may be mosquito physiology. Indeed, we note reports of bacterial community structures specific to mosquito organs such as midgut, salivary glands, ovaries, and Malpighian tubules, suggesting these tissues may reflect distinct niches in the mosquito [46,79–81]. One potential physiological difference that may differ between the crop and midgut is pH. The adult midgut has been shown to be actively maintained at slightly acidic conditions in sugar fed females (approximately pH 6.0) while the crop does not regulate the pH of its contents [82]. Therefore, if acid-producing bacteria colonize the crop, it may result in acidification which could influence growth of bacteria that cannot tolerate low pH. Gusmão et al. [20] found a dominant *Serratia* sp. growing in crops with the ability to produce acid when in the presence of glucose. In parallel, they observed acidification of crop contents over the course of 24 hours. In our study, *Tanticharoenia*, which is abundant in the crop, has the capacity to secrete acetic acid during fermentation. Notably, fermentation is an enriched functional process in the crop-associated bacterial community (Figure 8). We did not measure pH of the crop or midgut in our study; nonetheless, it is not unreasonable to hypothesize that AAB could be influencing bacterial diversity in the crop by altering the pH, while in the midgut, active regulation of the pH could limit the impact of acid producing bacteria and result in a different outcome in the community assembly process.

Microbial community structures may also be determined by the mosquitoes’ organ-specific immune responses. In the midgut, immune system signaling has been shown to control proliferation of bacteria [83–85]. Homeostasis within this complex system is regulated in part by a mechanism involving mosquito C-type lectins used by commensal bacteria to evade the effects of antimicrobial peptides (AMPs) [86,87]. To the best of our knowledge, the first study characterizing immune gene transcript abundance in the *Ae. aegypti* crop was recently published by Hixson et al. [88]. The authors propose that the predominant phenomenon in the crop would be immune recognition via orthologs of the Toll pathway. Similar to the posterior and hind gut, low abundance of gambicin and lyzozyme transcripts were detected in the crop. Segments with this transcriptional profile are hypothesized by the authors to be areas of microbial tolerance rather than bacterial community modulation as would be predicted in tissues with high AMP transcript abundance such as the midgut.

It is also possible that the differences in bacterial community structures between the crop and midgut were driven by bacteria-bacteria interactions within each niche. Bacteria interact metabolically with neighboring community members within short ranges [89]. The presence of a single nutrient source (e.g. carbohydrates) in a microenvironment can lead to bacterial exclusion and communities that are uneven and have low diversity [90]. This community dynamics model provides a plausible explanation for what we observed in our study. Alternatively, the dominance of *Elizabethkingia* in midguts could derive from antagonistic bacteria-bacteria interactions. It was recently reported that when co-cultured, *Elizabethkingia anophelis* inhibits the growth of a *Pseudomonas* sp. through an antimicrobial-independent mechanism [91]. In addition, the genome of *Elizabethkingia meningoseptica* contains genes predicted to confer resistance to multiple antibiotics [91–93] and other toxic compounds [93], potentially conferring advantage over other commensals when in hostile environments.

Many of the dominant bacteria in the crops (namely the Acetobacteraceae) were also found in the midgut tissues, albeit in smaller proportions and amounts than in the crops. This may indicate that some bacteria are transferred between these tissues. An example of this may occur if bacteria accompany the sugar that is pumped from the crop into the midgut when the mosquito needs to absorb its sugar reserves. However, it is possible that the proventriculus prevents most bacteria from moving from the foregut into the midgut as is seen with other insects [94]. Lanan et al. [94] demonstrated that the proventriculus acts as a bacterial filter and has a similar bacterial structure to the crop. Because our midgut samples included the proventriculus, the small amounts of crop-associated bacteria found in midguts may actually represent bacteria existing in the proventriculus. As bacteria transit from the mosquito crop to the midgut, gambicin and lysozyme produced in the proventriculus may exert selective pressure. In this scenario, the host’s immune system would be shaping the seeding community of the posterior gut segments [88].

Spatiotemporal dynamics may also be a critical determinant of community assembly along the digestive tract. As colonizing microbes enter the community either through internal transstadial transmission or ingestion of water and/or nectar [6,8,9,95], the invading order of each species would determine the community structure at each habitat. This phenomenon, known as the multiple stable states [96,97] has been tested in a *Drosophila* gut model [98]. This study revealed how, as each habitat along the gut faces the influx of new species, stochasticity and bottlenecks induce (probabilistic) hysteresis in the colonization process. Though deterministic processes (mediated by tissue attachment) could be at work, both crop and midgut communities would be unstable habitats in the fly model as they are colonized by bacteria in the lumen [98]. Studies under this ecological framework are yet to be performed in mosquito models, though the relevance of community ecology approaches has been raised to understand mosquito-microbiome interactions [49].

No functional studies have been performed, to the best of our knowledge, characterizing the role bacteria may play in the crop’s metabolic pathways. David et al. [62] report a functional analysis of midgut samples from sugarfed females using PICRUST. Though the functional categories are different to those in FAPROTAX, both approaches suggest the potential of the midgut bacterial community to perform carbon, nitrogen, and energy metabolism. Consistently with other models [99,100], we observed how the functional trait abundances of the bacterial communities reflect the overall beta diversity, as samples cluster predominantly in a by-tissue pattern. We detected a significant difference in methanol oxidation potential between the crop and midgut. Methanol catabolizing bacteria are methylotrophs and methanotrophs that establish and thrive when low pH acts as a niche-defining factor [101,102]. These organisms develop in the presence of plant derived sugars, being predominant members of the Earth’s phyllosphere [101,102]. The community enriched functions we report could be attributed to crop exclusive methylotrophs like *Methylobacterium* and *Methylovirgula* present in low abundances in the bacterial assembly. While we do not know whether methanol detoxification is a relevant phenomenon for adult mosquitos, it has been reported to be for mosquito larvae [103] and other insects that feed on plants [104].

Given that commensal community structures are influenced by deterministic and neutral processes when colonizing a niche along an insect’s gut, spatiotemporal scales and hysteresis in the community assembly process have to be considered [98]. As such, we acknowledge that the community structures we report for each organ are community snapshots given particular physiological host states, contextualized by the sex, age and nutritional states of the mosquitos we used. Nevertheless, our study demonstrates that different sections of the mosquito digestive tract can contain dramatically different communities of bacteria. Most notably, our study showed that the mosquito midgut is often overwhelmingly inhabited by *Elizabethkingia*, while the crop is generally more diverse and has a higher proportion of *Asaia* and *Tanticharoenia* bacteria. These distinct bacterial profiles may confer different biological functions, such as altering pH or metabolizing particular compounds. It is likely that multiple factors influence the development of these communities, such as host intrinsic parameters, bacteria-bacteria interactions, exchange of bacteria between tissues, and stochastic effects. Understanding how these communities assemble may facilitate better methods to engineer a mosquito microbiome that is refractory to pathogen transmission with the ultimate goal of reducing the disease burden that mosquitoes, especially *Ae. aegypti*, pose to people around the world [46,49, 105]. In addition, future work investigating organ-specific microbial communities has the potential to elucidate how production and exchange of secondary metabolites can influence within-host bacterial community dynamics and host health as seen in other models [106].

## Supporting information

Supplementary File 1_Raw data

16S profiling R script

## Data availability

Raw data is contained within Supplementary File 1, with the exception of raw sequence reads from high throughput amplicon sequencing. These sequences will be uploaded to the Sequence Read Archive upon acceptance.

## Acknowledgments

We wish to acknowledge the insectary staff at the Johns Hopkins Bloomberg School of Public Health for assistance with mosquito rearing.

---

**Supplementary Figure 1:**
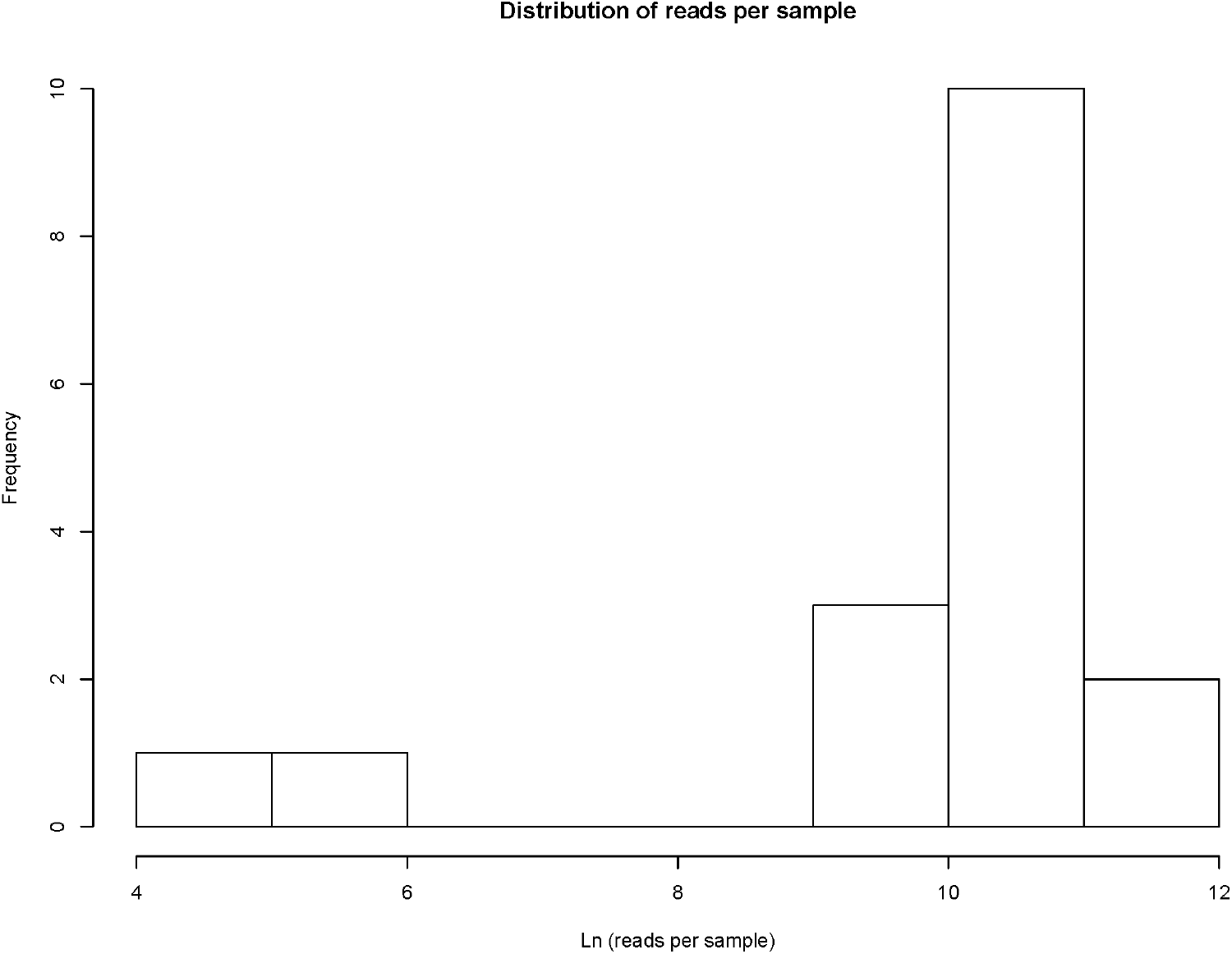
Histogram showing read counts for all samples.

**Supplementary Figure 2:**
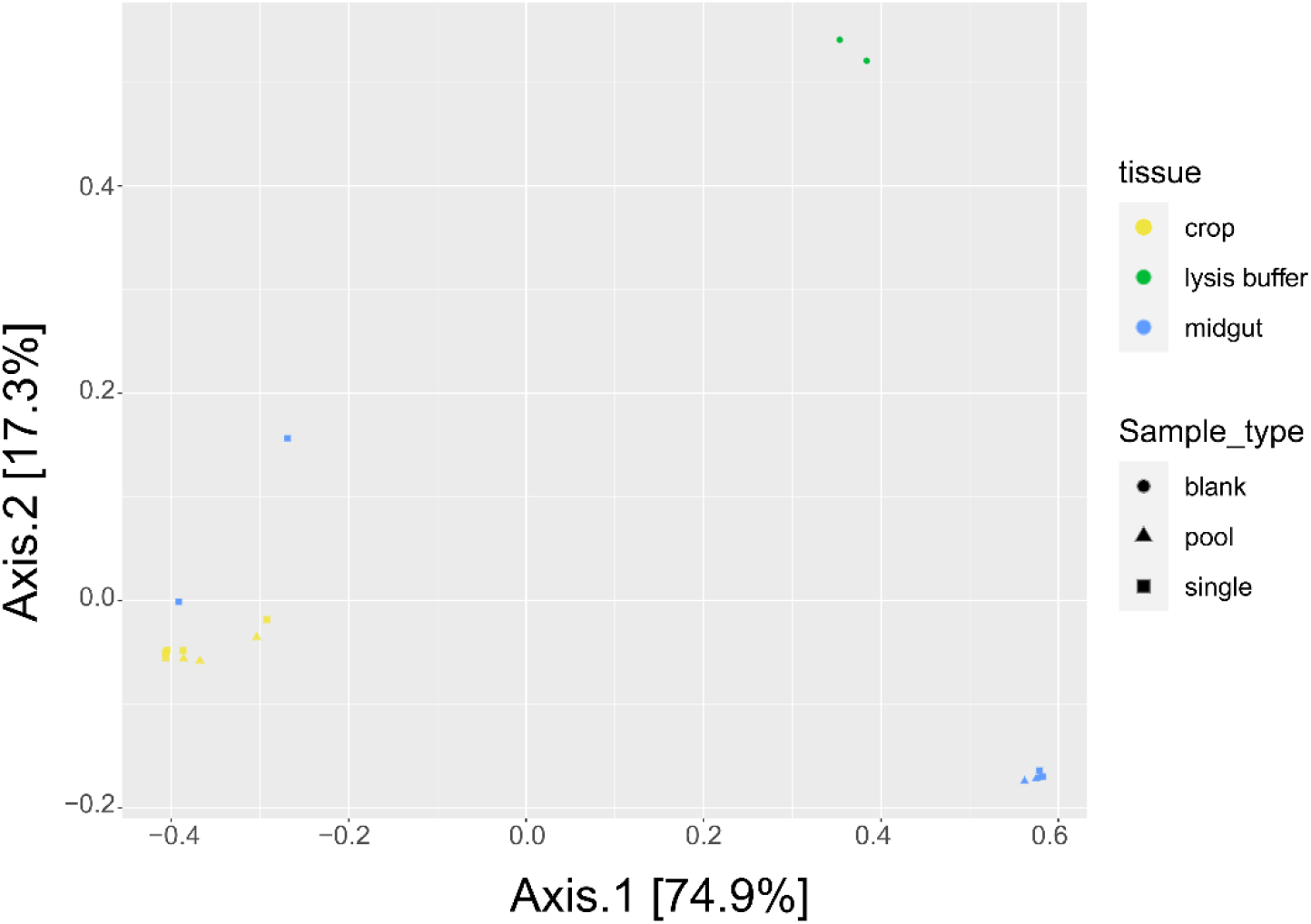
PCoA for all samples including buffer blanks. PCoA using Bray Curtis dissimilarity values shows that buffer blanks (shown in green) group together and separately from experimental samples.

## References

1. World Health Organization. Global vector control response 2017-2030 [Internet]. Geneva; 2017. Available from: http://apps.who.int

2. da Silveira LTC, Tura B, Santos M. Systematic review of dengue vaccine efficacy. BMC Infect Dis. 2019 Aug 28;19(1):750.

3. Liang H, Lee M, Jin X. Guiding dengue vaccine development using knowledge gained from the success of the yellow fever vaccine. Cell Mol Immunol. 2016 Jan;13(1):36–46.

4. Weaver SC. Prediction and prevention of urban arbovirus epidemics: a challenge for the global virology community. Antiviral Res. 2018 Aug 1;156:80–4.

5. Merritt RW, Dadd RH, Walker ED. Feeding behavior, natural food, and nutritional relationships of larval mosquitoes. Annu Rev Entomol. 1992 Jan;37(1):349–74.

6. Coon KL, Vogel KJ, Brown MR, Strand MR. Mosquitoes rely on their gut microbiota for development. Mol Ecol. 2014;23(11):2727–39.

7. Coon KL, Valzania L, McKinney DA, Vogel KJ, Brown MR, Strand MR. Bacteria-mediated hypoxia functions as a signal for mosquito development. Proc Natl Acad Sci USA. 2017;114(27):E5362–9.

8. Lindh JM, Borg-Karlson AK, Faye I. Transstadial and horizontal transfer of bacteria within a colony of *Anopheles gambiae* (Diptera: Culicidae) and oviposition response to bacteria-containing water. Acta Trop. 2008 Sep;107(3):242–50.

9. Moll RM, Romoser WS, Modrzakowski MC, Moncayo AC, Lerdthusnee K. Meconial peritrophic membranes and the fate of midgut bacteria during mosquito (Diptera: Culicidae) metamorphosis. J Med Entomol. 2001 Jan;38(1):29–32.

10. Saab SA, Dohna H zu, Nilsson LKJ, Onorati P, Nakhleh J, Terenius O, et al. The environment and species affect gut bacteria composition in laboratory co-cultured *Anopheles gambiae* and *Aedes albopictus* mosquitoes. Sci Rep. 2020 Feb 25;10(1):3352.

11. Caragata EP, Tikhe CV, Dimopoulos G. Curious entanglements: interactions between mosquitoes, their microbiota, and arboviruses. Curr Res Virol. 2019 Aug 1;37:26–36.

12. Gaio A de O, Gusmão DS, Santos AV, Berbert-Molina MA, Pimenta PF, Lemos FJ. Contribution of midgut bacteria to blood digestion and egg production in *Aedes aegypti* (Diptera: Culicidae) (L.). Parasit Vectors. 2011 Jun 14;4(1):105.

13. Xi Z, Ramirez JL, Dimopoulos G. The *Aedes aegypti* Toll pathway controls dengue virus infection. PLoS Pathog. 2008 Jul 4;4(7):e1000098.

14. Apte-Deshpande AD, Paingankar M, Gokhale MD, Deobagkar DN. *Serratia odorifera* a midgut inhabitant of *Aedes aegypti* mosquito enhances its susceptibility to dengue-2 virus. Vasilakis N, editor. PLoS ONE. 2012 Jul 27;7(7):e40401.

15. Apte-Deshpande AD, Paingankar MS, Gokhale MD, Deobagkar DN. *Serratia odorifera* mediated enhancement in susceptibility of *Aedes aegypti* for chikungunya virus. Indian J Med Res. 2014;139:762–8.

16. Ramirez JL, Short SM, Bahia AC, Saraiva RG, Dong Y, Kang S, et al. *Chromobacterium* Csp_P reduces malaria and dengue infection in vector mosquitoes and has entomopathogenic and in vitro anti-pathogen activities. Levashina E, editor. PLoS Pathog. 2014 Oct 23;10(10):e1004398.

17. Wu P, Sun P, Nie K, Zhu Y, Shi M, Xiao C, et al. A gut commensal bacterium promotes mosquito permissiveness to arboviruses. Cell Host Microbe. 2019 Jan 9;25(1):101–112.e5.

18. Jupatanakul N, Sim S, Dimopoulos G. The insect microbiome modulates vector competence for arboviruses. Viruses. 2014 Nov;6(11):4294–313.

19. Saraiva RG, Kang S, Simões ML, Angleró-Rodríguez YI, Dimopoulos G. Mosquito gut antiparasitic and antiviral immunity. Dev Comp Immunol. 2016 Nov 1;64:53–64.

20. Gusmão DS, Santos AV, Marini DC, Russo É de S, Peixoto AMD, Bacci Júnior M, et al. First isolation of microorganisms from the gut diverticulum of *Aedes aegypti* (Diptera: Culicidae): new perspectives for an insect-bacteria association. Mem Inst Oswaldo Cruz. 2007 Dec;102(8):919–24.

21. Megahed MM. Anatomy and histology of the alimentary tract of the female of the biting midge *Culicoides nubeculosus* Meigen (Diptera: Heleidae=Ceratopogonidae). Parasitol. 1956;46(1–2):22–47.

22. Cox JA. Morphology of the digestive tract of the blackfly (*Simulium nigroparvum*). J Agric Res. 1938;9.

23. Klowden MJ. Physiological systems in insects. 3rd ed. London: Academic Press; 2013.

24. Day MF. The mechanism of food distribution to the midgut or diverticula in the mosquito. Aust J Biol Sci. 1954;

25. Foster WA. Mosquito sugar feeding and reproductive energetics. Annu Rev Entomol. 1995;40:443–74.

26. Trembley HL. The distribution of certain liquids in the esophageal diverticula and stomach of mosquitoes. Am J Trop Med Hyg. 1952 Jul;1(4):693–710.

27. Calkins TL, DeLaat A, Piermarini PM. Physiological characterization and regulation of the contractile properties of the mosquito ventral diverticulum (crop). J Insect Physiol. 2017 Nov;103:98–106.

28. Stoffolano JG, Haselton AT. The adult Dipteran crop: a unique and overlooked organ. Annu Rev of Entomol. 2013;58(1):205–25.

29. Stoffolano JG. Fly foregut and transmission of microbes. In: Jurenka R, editor. Advances in Insect Physiology [Internet]. Academic Press; 2019 [cited 2022 Apr 18]. p. 27–95. Available from: https://www.sciencedirect.com/science/article/pii/S0065280619300207

30. Guégan M, Martin E, Valiente Moro C. Comparative analysis of the bacterial and fungal communities in the gut and the crop of *Aedes albopictus* mosquitoes: a preliminary study. Pathogens. 2020 Aug 1;9(8):E628.

31. Sim S, Jupatanakul N, Ramirez JL, Kang S, Romero-Vivas CM, Mohammed H, et al. Transcriptomic profiling of diverse *Aedes aegypti* strains reveals increased basal-level immune activation in dengue virus-refractory populations and identifies novel virus-vector molecular interactions. PLOS Negl Trop Dis. 2013 Jul 4;7(7):e2295.

32. Weisburg WG, Barns SM, Pelletier DA, Lane DJ. 16S ribosomal DNA amplification for phylogenetic study. J of Bacteriol. 1991 Jan;173(2):697–703.

33. Hall T. BioEdit: a user-friendly biological sequence alignment editor and analysis program for Windows 95/98/NT [Internet]. London: Information Retrieval Ltd.; 1999 [cited 2022 Aug 4]. Available from: https://bioedit.software.informer.com

34. Altschul SF, Gish W, Miller W, Myers EW, Lipman DJ. Basic local alignment search tool. J Mol Biol. 1990 Oct 5;215(3):403–10.

35. MacLeod HJ, Dimopoulos G, Short SM. Larval diet abundance influences size and composition of the midgut microbiota of *Aedes aegypti* mosquitoes. Front Microbol. 2021.

36. Nadkarni MA, Martin FE, Jacques NA, Hunter N 2002. Determination of bacterial load by real-time PCR using a broad-range (universal) probe and primers set. Microbiol. 2002;148(1):257–66.

37. Caporaso JG, Lauber CL, Walters WA, Berg-Lyons D, Lozupone CA, Turnbaugh PJ, et al. Global patterns of 16S rRNA diversity at a depth of millions of sequences per sample. Proc Natl Acad Sci. 2011 Mar 15;108(supplement_1):4516–22.

38. Kozich JJ, Westcott SL, Baxter NT, Highlander SK, Schloss PD. Development of a dual-index sequencing strategy and curation pipeline for analyzing amplicon sequence data on the MiSeq Illumina sequencing platform. Appl Environ Microbiol. 2013 Sep;79(17):5112–20.

39. R Core Team. R: a language and environment for statistical computing [Internet]. Vienna, Austria: R Foundation for Computing; 2021. Available from: https://www.R-project.org/

40. Callahan BJ, McMurdie PJ, Rosen MJ, Han AW, Johnson AJA, Holmes SP. DADA2: high-resolution sample inference from Illumina amplicon data. Nat Methods. 2016 Jul;13(7):581–3.

41. Callahan BJ. DADA2 pipeline tutorial (1.16) [Internet]. 2022 [cited 2022 Aug 7]. Available from: https://benjjneb.github.io/dada2/tutorial.html

42. Wang Q, Garrity GM, Tiedje JM, Cole JR. Naive Bayesian classifier for rapid assignment of rRNA sequences into the new bacterial taxonomy. Appl Environ Microbiol. 2007 Aug;73(16):5261–7.

43. McMurdie PJ, Holmes S. *phyloseq*: an R package for reproducible interactive analysis and graphics of microbiome census data. PLoS ONE. 2013 Apr 22;8(4):e61217.

44. Jervis-Bardy J, Leong LEX, Marri S, Smith RJ, Choo JM, Smith-Vaughan HC, et al. Deriving accurate microbiota profiles from human samples with low bacterial content through post-sequencing processing of Illumina MiSeq data. Microbiome. 2015 Dec;3(1):19.

45. Salter SJ, Cox MJ, Turek EM, Calus ST, Cookson WO, Moffatt MF, et al. Reagent and laboratory contamination can critically impact sequence-based microbiome analyses. BMC Biol. 2014 Dec;12(1):87.

46. Scolari F, Casiraghi M, Bonizzoni M. *Aedes* spp. and their microbiota: a review. Front Microbiol 2019;10.

47. Dada N, Jumas-Bilak E, Manguin S, Seidu R, Stenström TA, Overgaard HJ. Comparative assessment of the bacterial communities associated with *Aedes aegypti* larvae and water from domestic water storage containers. Parasit Vectors. 2014 Aug 24;7(1):391.

48. Hery L, Guidez A, Durand AA, Delannay C, Normandeau-Guimond J, Reynaud Y, et al. Natural variation in physicochemical profiles and bacterial communities associated with *Aedes aegypti* breeding sites and larvae on Guadeloupe and French Guiana. Microb Ecol. 2021 Jan;81(1):93–109.

49. Steven B, Hyde J, LaReau JC, Brackney DE. The axenic and gnotobiotic mosquito: emerging models for microbiome host interactions. Front Microbiol. 2021 Jul 12;12:714222.

50. Davis NM, Proctor DM, Holmes SP, Relman DA, Callahan BJ. Simple statistical identification and removal of contaminant sequences in marker-gene and metagenomics data. Bioinform; 2017.

51. Denef VJ, Fujimoto M, Berry MA, Schmidt ML. Seasonal succession leads to habitat-dependent differentiation in ribosomal RNA:DNA ratios among freshwater lake bacteria. Front Microbiol. 2016;7.

52. Fox J, Weisburg S. An R companion to applied regression [Internet]. Thousand Oaks, CA: Sage; 2019. Available from: https://socialsciences.mcmaster.ca/jfox/Books/Companion/

53. RStudio Team. RStudio: integrated development for R. [Internet]. Boston, MA: RStudio, PBC; 2020. Available from: http://www.rstudio.com/

54. Hill TCJ, Walsh KA, Harris JA, Moffett BF. Using ecological diversity measures with bacterial communities. FEMS Microbiol Ecol. 2003 Feb;43(1):1–11.

55. Kassambara A. ggpubr: “ggplot2” based publication ready plots [Internet]. 2020. Available from: https://rpkgs.datanovia.com/ggpubr/

56. Liu C, Cui Y, Li X, Yao M. *microeco* : an R package for data mining in microbial community ecology. FEMS Microbiol Ecol. 2021 Jan 26;97(2):fiaa255.

57. Oksanen J, Blanchet FG, Friendly M, Kindt R, Legendre P, McGlinn D, et al. vegan: community ecology package [Internet]. 2020. Available from: https://CRAN.R-project.org/package=vegan

58. Nearing JT, Douglas GM, Hayes MG, MacDonald J, Desai DK, Allward N, et al. Microbiome differential abundance methods produce different results across 38 datasets. Nat Commun. 2022 Jan 17;13(1):342.

59. Segata N, Izard J, Waldron L, Gevers D, Miropolsky L, Garrett WS, et al. Metagenomic biomarker discovery and explanation. Genome Biol. 2011 Jun 24;12(6):R60.

60. De Cáceres M, Legendre P. Associations between species and groups of sites: indices and statistical inference. Ecology. 2009;90(12):3566–74.

61. Louca S, Parfrey L, Doebeli M. Decoupling function and taxonomy in the global ocean microbiome. Science. 2016;353(6305):1272–7.

62. David MR, dos Santos LMB, Vicente ACP, Maciel-de-Freitas R. Effects of environment, dietary regime and ageing on the dengue vector microbiota: evidence of a core microbiota throughout *Aedes aegypti* lifespan. Mem Inst Oswaldo Cruz. 2016 Sep;111(9):577–87.

63. Muturi EJ, Ramirez JL, Rooney AP, Kim CH. Comparative analysis of gut microbiota of mosquito communities in central Illinois. PLOS Negl Trop Dis. 2017 Feb 28;11(2):e0005377.

64. Muturi EJ, Lagos-Kutz D, Dunlap C, Ramirez JL, Rooney AP, Hartman GL, et al. Mosquito microbiota cluster by host sampling location. Parasit Vectors. 2018 Aug 14;11(1):468.

65. Strand MR. The gut microbiota of mosquitoes: diversity and function. In: Wikel SK, Aksoy S, Dimopoulos G, editors. Arthropod Vector: Controller of Disease Transmission, Volume 1 [Internet]. Academic Press; 2017 [cited 2022 Aug 5]. p. 185–99. Available from: https://www.sciencedirect.com/science/article/pii/B9780128053508000118

66. Cao Q, Sun X, Rajesh K, Chalasani N, Gelow K, Katz B, et al. Effects of rare microbiome taxa filtering on statistical analysis. Front Microbiol. 2021;11.

67. Mamlouk D, Gullo M. Acetic acid bacteria: physiology and carbon sources oxidation. Indian J Microbiol. 2013 Dec 1;53(4):377–84.

68. Yukphan P, Malimas T, Muramatsu Y, Takahashi M, Kaneyasu M, Tanasupawat S, et al. *Tanticharoenia sakaeratensis* gen. nov., sp. nov., a new osmotolerant acetic acid bacterium in the alpha-Proteobacteria. Biosci Biotechnol Biochem. 2008 Mar;72(3):672–6.

69. Vu HTL, Malimas T, Chaipitakchonlatarn W, Bui VTT, Yukphan P, Bui UTT, et al. *Tanticharoenia aidae* sp. nov., for acetic acid bacteria isolated in Vietnam. Ann Microbiol. 2016 Mar;66(1):417–23.

70. Kawai M, Higashiura N, Hayasaki K, Okamoto N, Takami A, Hirakawa H, et al. Complete genome and gene expression analyses of *Asaia bogorensis* reveal unique responses to culture with mammalian cells as a potential opportunistic human pathogen. DNA Res. 2015 Oct;22(5):357–66.

71. Ramirez JL, Souza-Neto J, Torres Cosme R, Rovira J, Ortiz A, Pascale JM, et al. Reciprocal tripartite interactions between the *Aedes aegypti* midgut microbiota, innate immune system and dengue virus influences vector competence. PLoS Negl Trop Dis. 2012;6(3):e1561.

72. Audsley MD, Ye YH, McGraw EA. The microbiome composition of *Aedes aegypti* is not critical for *Wolbachia*-mediated inhibition of dengue virus. PLoS Negl Trop Dis. 2017 Mar 7;11(3):e0005426.

73. Coon KL, Brown MR, Strand MR. Mosquitoes host communities of bacteria that are essential for development but vary greatly between local habitats. Mol Ecol. 2016;25(22):5806–26.

74. Terenius O, Lindh JM, Eriksson-Gonzales K, Bussière L, Laugen AT, Bergquist H, et al. Midgut bacterial dynamics in *Aedes aegypti*. FEMS Microbiol Ecol. 2012 Jun 1;80(3):556–65.

75. Chen S, Blom J, Walker ED. Genomic, physiologic, and symbiotic characterization of *Serratia marcescens* strains isolated from the mosquito *Anopheles stephensi*. Front Microbiol. 2017;0.

76. Short SM, Mongodin EF, MacLeod HJ, Talyuli OAC, Dimopoulos G. Amino acid metabolic signaling influences *Aedes aegypti* midgut microbiome variability. PLOS Negl Trop Dis. 2017 Jul 28;11(7):e0005677.

77. Dickson LB, Ghozlane A, Volant S, Bouchier C, Ma L, Vega-Rúa A, et al. Diverse laboratory colonies of *Aedes aegypti* harbor the same adult midgut bacterial microbiome. Parasites & Vectors. 2018 Mar 27;11(1):207.

78. Schmitt A, Roy R, Carter CJ. Nectar antimicrobial compounds and their potential effects on pollinators. Curr Opin Insect Sci. 2021 Apr 1;44:55–63.

79. Díaz S, Camargo C, Avila FW. Characterization of the reproductive tract bacterial microbiota of virgin, mated, and blood-fed *Aedes aegypti* and *Aedes albopictus* females. Parasit Vectors. 2021 Dec 1;14(1):592.

80. Mancini MV, Damiani C, Accoti A, Tallarita M, Nunzi E, Cappelli A, et al. Estimating bacteria diversity in different organs of nine species of mosquito by next generation sequencing. BMC Microbiol. 2018 Oct 4;18:126.

81. Segata N, Baldini F, Pompon J, Garrett WS, Truong DT, Dabiré RK, et al. The reproductive tracts of two malaria vectors are populated by a core microbiome and by gender-and swarm-enriched microbial biomarkers. Sci Rep. 2016 Apr 18;6(1):24207.

82. Nepomuceno DB, Santos VC, Araújo RN, Pereira MH, Sant’Anna MR, Moreira LA, et al. pH control in the midgut of *Aedes aegypti* under different nutritional conditions. J Exp Biol. 2017 Sep 15;220(Pt 18):3355–62.

83. Clayton AM, Cirimotich CM, Dong Y, Dimopoulos G. Caudal is a negative regulator of the *Anopheles* IMD pathway that controls resistance to *Plasmodium falciparum* infection. Dev Comp Immunol. 2013 Apr;39(4):323–32.

84. Meister S, Agianian B, Turlure F, Relógio A, Morlais I, Kafatos FC, et al. *Anopheles gambiae* PGRPLC-mediated defense against bacteria modulates infections with malaria parasites. PLoS Pathog. 2009 Aug 7;5(8):e1000542.

85. Song X, Wang M, Dong L, Zhu H, Wang J. PGRP-LD mediates *A. stephensi* vector competency by regulating homeostasis of microbiota-induced peritrophic matrix synthesis. PLoS Pathog. 2018 Feb;14(2).

86. Li HH, Cai Y, Li JC, Su MP, Liu WL, Cheng L, et al. C-type lectins link immunological and reproductive processes in *Aedes aegypti*. iScience. 2020 Sep 25;23(9):101486.

87. Pang X, Xiao X, Liu Y, Zhang R, Liu J, Liu Q, et al. Mosquito C-type lectins maintain gut microbiome homeostasis. Nat Microbiol. 2016 Mar 14;1(5):1–11.

88. Hixson B, Bing XL, Yang X, Bonfini A, Nagy P, Buchon N. A transcriptomic atlas of *Aedes aegypti* reveals detailed functional organization of major body parts and gut regional specializations in sugar-fed and blood-fed adult females. Lemaître B, Banerjee U, Pondeville E, editors. eLife. 2022 Apr 26;11:e76132.

89. Dal Co A, van Vliet S, Kiviet DJ, Schlegel S, Ackermann M. Short-range interactions govern the dynamics and functions of microbial communities. Nat Ecol Evol. 2020 Mar;4(3):366–75.

90. Ratzke C, Barrere J, Gore J. Strength of species interactions determines biodiversity and stability in microbial communities. Nat Ecol Evol. 2020 Mar;4(3):376–83.

91. Ganley JG, D’Ambrosio HK, Shieh M, Derbyshire ER. Coculturing of mosquito-microbiome bacteria promotes heme degradation in *Elizabethkingia anophelis*. Chembiochem. 2020 May 4;21(9):1279–84.

92. Hyde J, Gorham C, Brackney DE, Steven B. Antibiotic resistant bacteria and commensal fungi are common and conserved in the mosquito microbiome. PLOS ONE. 2019 Aug 14;14(8):e0218907.

93. Kukutla P, Lindberg BG, Pei D, Rayl M, Yu W, Steritz M, et al. Draft genome sequences of *Elizabethkingia anophelis* strains R26T and Ag1 from the midgut of the malaria mosquito *Anopheles gambiae*. Genome Announc. 2013 Dec 5;1(6):e01030–13.

94. Lanan MC, Rodrigues PAP, Agellon A, Jansma P, Wheeler DE. A bacterial filter protects and structures the gut microbiome of an insect. ISME J. 2016 Aug;10(8):1866–76.

95. Bassene H, Niang EHA, Fenollar F, Doucoure S, Faye O, Raoult D, et al. Role of plants in the transmission of *Asaia* sp., which potentially inhibit the *Plasmodium* sporogenic cycle in *Anopheles* mosquitoes. Sci Rep. 2020 Apr 28;10(1):7144.

96. Abramsky Z, Rosenzweig ML. Tilman’s predicted productivity–diversity relationship shown by desert rodents. Nature. 1984 May;309(5964):150–1.

97. Goyal A, Dubinkina V, Maslov S. Multiple stable states in microbial communities explained by the stable marriage problem. ISME J. 2018 Dec;12(12):2823–34.

98. Obadia B, Güvener ZT, Zhang V, Ceja-Navarro JA, Brodie EL, Ja WW, et al. Probabilistic invasion underlies natural gut microbiome stability. Curr Biol. 2017 Jul 10;27(13):1999–2006.e8.

99. Fierer N, Leff JW, Adams BJ, Nielsen UN, Bates ST, Lauber CL, et al. Cross-biome metagenomic analyses of soil microbial communities and their functional attributes. Proc Natl Acad Sci U S A. 2012 Dec 26;109(52):21390–5.

100. Ruhl IA, Sheremet A, Smirnova AV, Sharp CE, Grasby SE, Strous M, et al. Microbial functional diversity correlates with species diversity along a temperature gradient. mSystems. 2022 Feb 15;7(1):e00991–21.

101. Stacheter A, Noll M, Lee CK, Selzer M, Glowik B, Ebertsch L, et al. Methanol oxidation by temperate soils and environmental determinants of associated methylotrophs. ISME J. 2013 May;7(5):1051–64.

102. Morawe M, Hoeke H, Wissenbach DK, Lentendu G, Wubet T, Kröber E, et al. Acidotolerant bacteria and fungi as a sink of methanol-derived carbon in a deciduous forest soil. Front Microbiol. 2017 Jul 24;8:1361.

103. Liu XH, Pan H, Mazur P. Permeation and toxicity of ethylene glycol and methanol in larvae of *Anopheles gambiae*. J Exp Biol. 2003 Jul;206(Pt 13):2221–8.

104. Dixit S, Upadhyay SK, Singh H, Sidhu OP, Verma PC, K C. Enhanced methanol production in plants provides broad spectrum insect resistance. PLoS One. 2013;8(11):e79664.

105. Guégan M, Zouache K, Démichel C, Minard G, Tran Van V, Potier P, et al. The mosquito holobiont: fresh insight into mosquito-microbiota interactions. Microbiome. 2018 Mar 20;6(1):49.

106. Martínez JE, Vargas A, Pérez-Sánchez T, Encío IJ, Cabello-Olmo M, Barajas M. Human microbiota network: unveiling potential crosstalk between the different microbiota ecosystems and their role in health and disease. Nutrients. 2021 Aug 24;13(9):2905.

