## Supplementary material for "Bacterial communities of *Aedes aegypti* mosquitoes differ between crop and midgut tissues": 16S profiling R script

## Crop and Midgut Microbiome 16S profiling
# Target region: hypervariable region V4

# Analysis performed the servers made avaialbe by Fiocruz MG. Brazil.We acknowledge the 
# courtesy of Fausto Goncalves and Gabriel Fernandes for providing access and support to carry this analysis.


# Setting the working directory where the raw files are located.
setwd("~/MidgutCrop")

## Bioinformatic analyses performed following dada2 workflow v1.16 at: https://benjjneb.github.io/dada2/tutorial.html

# Load package
library(dada2); packageVersion("dada2")

#Specify path where FASTQ files are located
pathMC <- "/home/lemv/MidgutCrop"
pathMC
list.files(pathMC)

#Sort files to ensure forward/reverse reads are in the same order
# Forward and reverse fastq filenames have format: SAMPLENAME_L001_R1_001.fastq and SAMPLENAME_R2_001.fastq
fnFsMC <- sort(list.files(pathMC, pattern = "_L001_R1_001.fastq.gz",full.names = TRUE))
fnFsMC
fnRsMC <- sort(list.files(pathMC, pattern = "_L001_R2_001.fastq.gz",full.names = TRUE))
fnRsMC
basename(fnFsMC)

# Extract sample names, assuming filenames have format: SAMPLENAME_XXX.fastq
sample.namesMC <- sapply(strsplit(basename(fnFsMC), "_"), `[`, 1)
sample.namesMC

#Inspect read quality profiles
plotQualityProfile(fnFsMC[1:2])
plotQualityProfile(fnFsMC[9:10])
plotQualityProfile(fnFsMC[16:17]) ## considerably less reads: 396 and 238 for lysis buffer samples
# do this for samples and the blank

# Now the reverse reads
plotQualityProfile(fnRsMC[1:2])
plotQualityProfile(fnRsMC[9:10])
plotQualityProfile(fnRsMC[16:17])

# Place filtered files in filtered/ subdirectory
filtFsMC <- file.path(pathMC, "filtered", paste0(sample.namesMC, "_F_filt.fastq.gz"))
filtRsMC <- file.path(pathMC, "filtered", paste0(sample.namesMC, "_R_filt.fastq.gz"))
names(filtFsMC) <- sample.namesMC
names(filtRsMC) <- sample.namesMC

## Filter and trim the sequences as suggested for V4 amplicons
outMC <- filterAndTrim(fnFsMC, filtFsMC, fnRsMC, filtRsMC, truncLen=c(240,160),
                       maxN=0, maxEE=c(2,2), truncQ=2, rm.phix=TRUE,
                       compress=TRUE, multithread=TRUE)
head(outMC)

# Learning the error rates
errF <- learnErrors(filtFsMC, multithread=TRUE)
errR <- learnErrors(filtRsMC, multithread=TRUE)

# Applying sample inference algorithm.

# Extensions: By default, the dada function processes each sample independently. 
# However, pooling information across samples can increase sensitivity to sequence variants that may be present at very low frequencies in multiple samples. 
# We used the standard pooling alternative (..., pool=TRUE) in which all samples are pooled together for sample inference. 

dadaFs <- dada(filtFsMC, err=errF, multithread=TRUE,pool=TRUE)
dadaRs <- dada(filtRsMC, err=errR, multithread=TRUE, pool=TRUE)

dadaFs[[1]]

# Forward and reverse reads merging
mergers <- mergePairs(dadaFs, filtFsMC, dadaRs, filtRsMC, verbose=TRUE)
# Inspect the merger data.frame from the first sample
head(mergers[[1]])

# Amplicon sequence variant table construction
seqtab <- makeSequenceTable(mergers)
dim(seqtab)

# Inspect distribution of sequence lengths
table(nchar(getSequences(seqtab)))

# Chimera removal
seqtab.nochim <- removeBimeraDenovo(seqtab, method="consensus", multithread=TRUE, verbose=TRUE)
dim(seqtab.nochim)
# [1]  17 289  --> 289 ASVs are true non chimeric ASVs
sum(seqtab.nochim)/sum(seqtab)
# 0.995272 --> chimeras account for ~1% of the merged sequences

# Tracking the reads as they go through the pipeline
getN <- function(x) sum(getUniques(x))
track <- cbind(outMC, sapply(dadaFs, getN), sapply(dadaRs, getN), sapply(mergers, getN), rowSums(seqtab.nochim))
# If processing a single sample, remove the sapply calls: e.g. replace sapply(dadaFs, getN) with getN(dadaFs)
colnames(track) <- c("input", "filtered", "denoisedF", "denoisedR", "merged", "nonchim")
rownames(track) <- sample.namesMC
head(track)

tail(track) ## blank samples
#                        input filtered denoisedF denoisedR merged nonchim
# SMSSeq2amplicon16SH09   396      303       290       296    255     255
# SMSSeq2amplicon16SH10   238      175       172       169    142     142


# Assigning the taxonomy 
# We downloaded the RDP training set v18 formatted for dada2 and available at: https://zenodo.org/record/4310151#.YoPmRWDMLKo 
taxa <- assignTaxonomy(seqtab.nochim, "~/Downloads/rdp_train_set_18.fa.gz", multithread=TRUE)

# species level assignment based on exact matching
taxa <- addSpecies(taxa, "~/Downloads/rdp_species_assignment_18.fa.gz")

# Inspecting the taxonimic assignments
taxa.print <- taxa # Removing sequence rownames for display only
rownames(taxa.print) <- NULL
head(taxa.print)
tail(taxa.print)

#######   PHYLOSEQ   ###########

# Importing the dada2 output (count table and taxonomy) to Phyloseq
# Load required packages
library(phyloseq); packageVersion("phyloseq")
library(Biostrings); packageVersion("Biostrings")
library(ggplot2); packageVersion("ggplot2")

# Importing sample metadata including: sample name, tissue of origin, sample type (pool or single), sample ID, etc.
SampledataMC<-read.csv("~/Downloads/MC_SampleData.txt", header = TRUE, sep = ";", row.names=1)
SampledataMC
class(SampledataMC)


# creating Phyloseq object: OTU table, Tax table, Sample ID, Phylogeny, ref seq
#Extracting the standard goods from R
# giving the seq headers more manageable names (ASV_1, ASV_2...)
asv_seqsMC <- colnames(seqtab.nochim)
asv_seqsMC

# The dimensions of our matrix are 17 x 289
dim(seqtab.nochim)

asv_headersMC <- vector(dim(seqtab.nochim)[2], mode="character")  # vector with spaces corresponding to ASV names
asv_headersMC

for (i in 1:dim(seqtab.nochim)[2]) {
  asv_headersMC[i] <- paste(">ASV", i, sep="_")
}

asv_headersMC  # Vector with labels for the true ASVs detected, whose sequences are on asv_seqsT variable

# making and writing out a fasta of the final ASV seqs:
asv_fastaMC <- c(rbind(asv_headersMC, asv_seqsMC))
asv_fastaMC
write(asv_fastaMC, "ASVs_Midgut_crop_microbiota.fa")  #fasta file of each ASV for other analyses 

# count table:
asv_tabMC <- t(seqtab.nochim)
asv_tabMC
row.names(asv_tabMC) <- sub(">", "", asv_headersMC)
write.table(asv_tabMC, "ASVs_countsMidgut_crop.txt", sep="\t", quote=T)
asv_tabMC

ASVcountsMC <-t(asv_tabMC)
ASVcountsMC  # transposed and with ASV_ structure to build the phyloseq objetc compatible with our explanatory matrix: 
             # samples as rows.

# tax table:
asv_taxMC <- taxa
row.names(asv_taxMC) <- sub(">", "", asv_headersMC)
asv_taxMC
write.table(asv_taxMC, "ASVs_taxonomyMidgut_crop.txt", sep="\t", quote=T)

## PHYLOSEQ object

# assigning the same sample names to our metadata table
rownames(SampledataMC)<-rownames(ASVcountsMC)
rownames(SampledataMC)
SampledataMC

MCPS <- phyloseq(otu_table(ASVcountsMC, taxa_are_rows=FALSE), 
                   sample_data(SampledataMC), 
                   tax_table(asv_taxMC))
MCPS # observe the elements contained in the phyloseq object

# exploring the phyloseq object (e.g most abundant ASV tax ID)
tax_table(MCPS)[1,]

# Taxonomy Table:     [1 taxa by 7 taxonomic ranks]:
#        Kingdom    Phylum           Class                 Order              Family            
#ASV_1 "Bacteria" "Proteobacteria" "Alphaproteobacteria" "Rhodospirillales" "Acetobacteraceae"
#       Genus           Species
#ASV_1 "Tanticharoenia" NA     

# Filtering out spurious tax id calls like chloroplast and mitochondria
# Observing how many of the ID's are True to the statement tested. Column 3 is Class and column 5 is Family
tax_table(MCPS)[,3]=="Chloroplast"
tax_table(MCPS)[,5]=="Mitochondria"
  
# MCPS phyloseq object has 289 ASVs. We need to remove those whose tax ID is deemed as spurious in the community
MCPS <- subset_taxa(MCPS, (
  Kingdom=="Bacteria" &
    Family != "Mitochondria" &
    Class != "Chloroplast")
)

MCPS # the dataset now has 147 ASVs

# Stacked barplot of relative abundances at the family rank to visualize the composition of our samples and controls before QC
install.packages("circlize") # package that allows us to create a random colors vector
genlcolorMC<-rand_color(50,hue=NULL,luminosity="random",transparency =0)
genlcolorMC

# Proportions phyloseq object
MCPSprop<-transform_sample_counts(MCPS, function(x) x / sum(x) )
sample_sums(MCPSprop) # all sum 1
MCPSprop

# Creating a data frame to build a by sample proportions barplot
MCPSfammelt <- psmelt(MCPSprop) # create dataframe from phyloseq object
MCPSfammelt$Family <- as.character(MCPSfammelt$Family) #convert to character
MCPSfammelt$Family
head(MCPSfammelt) # Inspecting the DF
tail(MCPSfammelt)

#simple way to rename genera with < 1% abundance
MCPSfammelt$Family[MCPSfammelt$Abundance < 0.01] <- "< 1% abund."

# barplot element
barplotMCPSfam <- ggplot(data=MCPSfammelt, aes(x=Sample, y=Abundance, fill=Family))

# colors to be used as taxonomy legend
colorMCPS2<-rand_color(30,hue=NULL,luminosity = "random",transparency = 0.4)
colorMCPS2

# barplot with aesthetics and facets
barplotMCPSfam + geom_bar(aes(), stat="identity", position="stack") +
  scale_fill_manual(values = colorMCPS2) +
  theme(legend.position="bottom",axis.text.x = element_text(size=2,face="bold")) + guides(fill=guide_legend(nrow=5))+
  facet_grid(~tissue, scales = "free_x" )


#### Ordinations based on proportions data
sample_data(MCPSprop) # recalling the metadata to verify the name of the variables to use as explanatory to assigns color and shape

ordpropMCPS2<-ordinate(MCPSprop,"PCoA","bray")
plot_ordination(MCPSprop,ordpropMCPS2, type="samples", color="tissue",shape="Sample_type",
                title="Crop, midgut and buffer control composition") + geom_point(size=2,alpha=0.4)

   # NO clustering between experimental samples and controls. #

# Total number of individuals observed from each ASV
sum.check <- taxa_sums(MCPS)
sum(sum.check)#534462

# Other descriptive statistics:
print(sample_sums(MCPS)) 
median(sample_sums(MCPS))
## median 36343

# Control samples raw read abundance
#SMSSeq2amplicon16SH09 
#240

#SMSSeq2amplicon16SH10 
#133 

# Average
# (240+133)/2 = 186.5 
# Percentage of reads that could be potentially contaminants from the averages

#186.5/36343 = 0.005  --> 0.5% of reads are potentially contamination. From the ordination analysis we know these 2 samples 
# culster separately from the experimental samples. 

# Histogram to evaluate the break betwwen the experimental samples and controls based on their raw reads 
hist(log(sample_sums(MCPS)),breaks="sturges", main="Distribution of reads per sample", xlab="Ln (reads per sample)")


### DECONTAM:  removing contaminants by prevalence, using the blank samples sequenced

sample_data(MCPS)
sample_data(MCPS)$is.neg <- sample_data(MCPS)$Sample_type == "blank"
sample_data(MCPS)$is.neg
contamdf.prevMC <- isContaminant(MCPS, method="prevalence", neg="is.neg")
table(contamdf.prevMC$contaminant)
# FALSE  TRUE 
# 146    1 

contamdf.prevMC$contaminant
head(which(contamdf.prevMC$contaminant))
# [1]   141  --> ID of the cotaminant ASV in the contam table

# we have only 2 blank samples out of 17 ~11% of samples. We establish the threshold at 0.2 arbitrarily to still be stringent given the fact we only have these 2 blanks

contamdf.prev02MC <- isContaminant(MCPS, method="prevalence", neg="is.neg", threshold=0.2)
table(contamdf.prev05MC$contaminant)

#  FALSE  TRUE 
#  135    12   --> there were 11 more ASVs that can be deemed as contaminants as they are more prevalent in the blanks than in the samples
which(contamdf.prev02MC$contaminant)


MCPS # removing the 12 contaminant ASVs from the 147 in the dataset
MCPS_2 <- prune_taxa(!contamdf.prev02MC$contaminant, MCPS)
MCPS_2  # 135 taxa in the otu_table element as expected sans contaminants


## Verifying if we have singleton ASVs and samples with just as singleton 
sum(sample_sums(MCPS_2)==0) ## 0 with just a singleton, 0 with zero
sum(taxa_sums(MCPS_2)==0) # 0 ASVs with just one read represented in the table 


tax_table(MCPS_2)[,2]=="NA"  # we dont have ASVs that are Bacteria but NA at the Phylum level


# Removing the blanks from the data set by extracting the subset that belongs to samples type only
sample_data(MCPS_2)
# Type = Sample
MCPS_3b<-subset_samples(MCPS_2,Sample_type!="blank")
MCPS_3b

# Filtering out taxa with less than 5 reads (in total) as an additional rare feature removal startegy
MCPS_4A<-phyloseq_filter_prevalence(MCPS_3b,abund.trh = 5, abund.type = "total") 
MCPS_4A # we kept the 135 ASVs

# Exploring the data so far

# Dominant ASV per sample at genus level after decontam and filtering
dominant(MCPS_4A, level = "Genus")


# Prevalence plot at family level
phyloseq_prevalence_plot(MCPS_4A, prev.trh = 0.5, taxcolor = "Family",
                         facet = T, point_alpha = 0.7, showplot = T)


# Phyloseq subsets accounting for the dominant genera and to obtain raw counts to be used dowstream in CCA analysis

#Elizabethkingia
Elizabethkingia<-otu_table(MCPS_4A)[,2]
Elizabethkingia<-as.data.frame(Elizabethkingia)
sampleID<-sample_data(MCPS)$sampleID
rownames(Elizabethkingia)<-sampleID
colnames(Elizabethkingia)<-c("Etot")
Elizabethkingia$Total<-sample_sums(MCPS_4A)
Elizabethkingia$Ratio<-Elizabethkingia$Etot/Elizabethkingia$Total
Elizabethkingia$E_LOG10<-log10(Elizabethkingia$Etot+1)
Elizabethkingia$samptype<-sample_data(MCPS_4A)$Sample_type
Elizabethkingia$tissue<-sample_data(MCPS_4A)$tissue
dim(Elizabethkingia)
Elizabethkingia2<-Elizabethkingia[1:15,]
Elizabethkingia2

#Tanticharoenia
Tanticharoenia = subset_taxa(MCPS_4A,Genus %in% c("Tanticharoenia"))
Tanticharoenia
otu_table(Tanticharoenia)
TantTot<-sample_sums(Tanticharoenia)
TantTot<-as.data.frame(TantTot)
rownames(TantTot)<-sample_data(Tanticharoenia)$sampleID
TantTot[1:15,1]
TantTot2<-TantTot[1:15,1]
TantTot2<-as.data.frame(TantTot2)
rownames(TantTot2)<-sample_data(MCPS_4A)$sampleID
TantTot2

## Asaia
Asaia = subset_taxa(MCPS_4A,Genus %in% c("Asaia"))
Asaia
sample_data(Asaia)$sampleID
otu_table(Asaia)
AsaTot<-sample_sums(Asaia)
AsaTot<-as.data.frame(AsaTot)
rownames(AsaTot)<-sample_data(Asaia)$sampleID
AsaTot

AsaTot2<-AsaTot[1:15,1]
AsaTot2<-as.data.frame(AsaTot2)
rownames(AsaTot2)<-sample_data(MCPS_4A)$sampleID
AsaTot2

### Summaryzing the data set so far
summary(tax_table(MCPS_4A))


# Creating a data frame to generate relative abundance barplots at the family taxonomic level

##Color palette contrast. Looking for alternatives to make a bars with good contrast
## website https://medialab.github.io/iwanthue/ I want Hues package
install.packages('hues')
colorMCPS_4B<-iwanthue(18)  #rather than random, this are optically distinct colors
colorMCPS_4B

######
# Normalizing by depth of sequencing to address the potential disparities introduced by the 2 separate seq events

scale_reads <- function(physeq, n) {
  physeq.scale <-
    transform_sample_counts(physeq, function(x) {
      (n * x/sum(x))
    })
  otu_table(physeq.scale) <- floor(otu_table(physeq.scale))
  physeq.scale <- prune_taxa(taxa_sums(physeq.scale) > 0, physeq.scale)
  return(physeq.scale)
}

# We then subsample reads to even depth (lowest read count in dataset) using the ???scale_reads function???
MCPS_4Anormalized <- scale_reads(MCPS_4A, n=min(sample_sums(MCPS_4A)))   ####*****

#Did it work? Looking at the distribution of read counts:
sum(sample_sums(MCPS_4Anormalized))  
#138512 

min(sample_sums(MCPS_4Anormalized))
# 9221

max(sample_sums(MCPS_4Anormalized))
# 9246

mean(sample_sums(MCPS_4Anormalized))
# 9234.1

## Creating a dataset with proportions
MCPS_4Anormalized
MCPSprop2Anorm<-transform_sample_counts(MCPS_4Anormalized, function(x) x / sum(x) )
sample_sums(MCPSprop2Anorm) # all sum 1

otu_table(MCPSprop2Anorm)[,2]
MCPS_BAR <- psmelt(MCPSprop2Anorm) # create dataframe from phyloseq object
MCPS_BAR$Family <- as.character(MCPS_BAR$Family) #convert to character
MCPS_BAR

#simple way to rename genera with < 1% abundance
MCPS_BAR$Family[MCPS_BAR$Abundance < 0.01] <- "< 1% abund."

barplotMCPS_BAR <- ggplot(data=MCPS_BAR, aes(x=sampleID, y=Abundance, fill=Family))

colorMCPS_4B

barplotMCPS_BAR + geom_bar(aes(), stat="identity", position="stack") +
  scale_fill_manual(values = colorMCPS_4B)+
  theme(legend.position="bottom",axis.text.x = element_text(size=7,face="bold")) + guides(fill=guide_legend(nrow=5))+
  facet_wrap(Sample_type~tissue, scales= "free_x", strip.position=c("left"))


### ALPHA DIVERSITY

# Alpha Diversity using microbiome package to obtain equity indices (Gini)
dominant(MCPS_4Anormalized,level="Family")
dominance(MCPS_4Anormalized)

alphaMCPS_normA<-estimate_richness(MCPS_4Anormalized,split=T,measures=c("Observed", "Shannon", "Simpson","Chao1","ACE"))
alphaMCPS_normA  


## Building an alpha div DF to explore the results
alphaMCPS_normA$tissue<-sample_data(MCPS_4A)$tissue
alphaMCPS_normA$Sample_type<-sample_data(MCPS_4A)$Sample_type
alphaMCPS_normA$rep<-sample_data(MCPS_4)$pair_number
row.names(alphaMCPS_normA)<-sample_data(MCPS_4A)$sampleID
alphaMCPS_normA$Etot<-Elizabethkingia2$Etot
alphaMCPS_normA$sample_ID<-sample_data(MCPS_4A)$sampleID
alphaMCPS_normA


# Ploting alpha diversity
Alpha_tissue_poolNORMA<-plot_richness(MCPS_4Anormalized, measures=c("Chao1", "Shannon","Simpson","Observed"),x="tissue", color="Sample_type")
Alpha_tissue_poolNORMA + geom_boxplot(data=Alpha_tissue_poolNORMA$data, aes(x=tissue, y=value, color=NULL), alpha=0.1)

# or

# Box plot facetted by sample type (sigle vs pool)
boxMCPSalpha <- ggboxplot(alphaMCPS_normA, x = "tissue", y = "Simpson",
                          fill = "tissue", palette = c("#1b9e77","#d95f02"),
                          add = "jitter",
                          facet.by = "Sample_type", short.panel.labs = T,bxp.errorbar=T,bxp.errorbar.width = 0.4)


boxMCPSalpha + stat_compare_means(method="kruskal.test", label = "p.format") + theme(legend.position="none") 


boxMCPSalpha2 <- ggboxplot(alphaMCPS_normA, x = "tissue", y = "Observed",
                          fill = "tissue", palette = c("#1b9e77","#d95f02"),
                          add = "jitter",
                          facet.by = "Sample_type", short.panel.labs = T,bxp.errorbar=T,bxp.errorbar.width = 0.4)


boxMCPSalpha2 + stat_compare_means(method="kruskal.test", label = "p.format") + theme(legend.position="none") 


# Subset DF by sample type to evaluate variance homogeneity for each alpha div value considering sample type
alphadivSingle<-filter(alphaMCPS_normA, Sample_type == "single") 
alphadivSingle

alphadivPool<-filter(alphaMCPS_normA, Sample_type == "pool") 
alphadivPool


leveneTest(Simpson ~ tissue, center="mean",data = alphadivSingle)
leveneTest(Observed ~ tissue, center="mean",data = alphadivSingle)

leveneTest(Simpson ~ tissue, center="mean",data = alphadivPool)
leveneTest(Observed ~ tissue, center="mean",data = alphadivPool)


#### Beta diversity


# Extracting the abundances matrix from the phyloseq object to run ordination analysis
MCPSprop2Anorm
ASVcountMCPSN<-otu_table(MCPSprop2Anorm)
ASVcountMCPS2N<-as.data.frame(ASVcountMCPSN)
ASVcountMCPS2N<-as.matrix(ASVcountMCPS2N)
ASVcountMCPS2N


# Labels for ordination and permanova analyses
labelsMCPS<-as.data.frame(sample_data(MCPS_4Anormalized)$tissue)
labelsMCPS
row.names(labelsMCPS)<-row.names(sample_data(MCPS_4Anormalized))
colnames(labelsMCPS)
colnames(labelsMCPS)<-c("Tissue")
colnames(labelsMCPS)

labelsMCPS3<-as.data.frame(sample_data(MCPS_4Anormalized)$Etot)
labelsMCPS3
colnames(labelsMCPS3)<-c("Etot")
labelsMCPSchi3
sample_data(MCPS_4Anormalized)
labelsMCPS4<-as.data.frame(sample_data(MCPS_4Anormalized)$Sample_type)
colnames(labelsMCPS4)<-c("Sample_type")
colnames(labelsMCPS4)
labelsMCPS4

labelsMCPS2<- labelsMCPS2 %>% mutate(labelsMCPS3$Etot)
labelsMCPS2
colnames(labelsMCPS2)[2]<-c("Etot")

labelsMCPS2<- labelsMCPS2 %>% mutate(labelsMCPS4$Sample_type)
labelsMCPS2
colnames(labelsMCPS2)[3]<-c("Sample_type")
labelsMCPS2

# Unconstrained ordination
### Microeco PCoA (Figure 5A in the main manuscript) is linked to the below permanova analysis ###

###Permanova reveals that sample type has no interaction nor significant effect on betadiversity
permMCPSANORM<- adonis2(ASVcountMCPS2N~Tissue*Sample_type,data= labelsMCPS2, permutations=999, method="bray", by= "terms",na.rm=T)
permMCPSANORM

# Constrained ordination
### CCA analisis to assess the effect of the most abundante ASVs as drivers of betadiversity
labelsMCPS2A<-labelsMCPS2
rownames(labelsMCPS2A)<-rownames(ASVcountMCPS2A)
rownames(labelsMCPS2A)
labelsMCPS2A
labelsMCPS3<-labelsMCPSchi2A
labelsMCPS3$Tanticharoenia<-TantTot2$TantTot2
labelsMCPS3$Asaia<-AsaTot2$AsaTot2
labelsMCPSCCA<-labelsMCPSchi3
colnames(labelsMCPSCCA)<-c("Tissue", "Elizabethkingia", "Sample_type", "Tanticharoenia","Asaia")
labelsMCPSCCA


ASVcountMCPS2A<-ASVcountMCPS2N
rownames(ASVcountMCPS2A)<-sample_data(MCPS_4Anormalized)$sampleID
rownames(ASVcountMCPS2A)


CCAMCPS2<-cca(ASVcountMCPS2A~Tissue+(Etot+Asaia+Tanticharoenia),data= labelsMCPSCCA, scaling=T)
summary(CCAMCPS2)
ordiplot(CCAMCPS2, scaling=2)

#### Permanova to test for the significance of the variables

permMCPS4<- adonis2(ASVcountMCPS2N~(Tissue+Etot+Asaia+Tanticharoenia)*Sample_type,data= labelsMCPSchi3, permutations=999, method="bray", by= "terms",na.rm=T)
permMCPS4 #--> Sample type does not interact significantly with any of the variables 

permMCPS5<- adonis2(ASVcountMCPS2N~(Etot+Asaia+Tanticharoenia)*Tissue,data= labelsMCPSchi3, permutations=999, method="bray", by= "terms",na.rm=T)
permMCPS5 # --> assessing the interaction between tissue type and the dominant ASVs
#

## Improving the CCA plot for the manuscript

CCAplot<-ordiplot(CCAMCPS2,choices=c(1,2), type = "n", display =c("wa", "cn","sp","lc","bp"),
                  scaling = 2,ylab="CCA2: 16.82%",xlab="CCA1: 21.20%",xlim = c(-3,1.5),ylim = c(-1.5,1.5) )

vectorcolorMCPS <- c(rep("#1b9e77",3),rep("#d95f02",3), rep("#1b9e77",5), rep("#d95f02",4)) # RGB options http://www.endmemo.com/program/R/color.php
vectorcolorMCPS

points(CCAMCPS2, display = "sites",pch=10, col=vectorcolorMCPS, scaling=2, cex=1)

text(CCAMCPS2, display="bp", col="black", scaling=2, arrow.mul = 1.3,cex=1)

points(CCAMCPS2, display="cn",cex = 2, pch=18,col="black", scaling=2)
text(CCAMCPS2, display="cn",cex = 1, offset=0.6, scaling = 2, pos=1)


legend("bottomleft",title="Tissue type",
       c("crop","midgut"),ncol=3,cex=0.85,
       fill=c("#1b9e77","#d95f02"))


# Indicator spcies analysis (load indiscpec package)
indicMCPS<-multipatt(as.data.frame(otu_table(MCPS_4Anormalized)),labelsMCPSchi3$Tissue,duleg=F,func="IndVal.g" ,control=how(nperm=999))
indicMCPS
summary(indicMCPS)
MCPSIndic<-tax_table(MCPS_4Anormalized)[c("ASV_4","ASV_71"),]
kable(MCPSIndic)      # print the table with the taxonomic information for the indicator ASVs                


# Microeco package (load microeco package)

microMCPSnorm<-phyloseq2meco(MCPS_4Anormalized)
tidy_taxonomy(microMCPSnorm$tax_table)
print(microMCPSnorm)
microMCPSnorm$tidy_dataset()
print(microMCPSnorm)

microMCPSnorm$sample_sums() %>% range
#1] 9221 9246


## Estimating the prportions per ASVS by sample 
microMCPSnorm$cal_abund()
microMCPSnorm$taxa_abund

# Beta diversity estimation for the PCoA ordination with Bray Curtis dissimilarities
microMCPSnorm$cal_betadiv(unifrac = FALSE)


BetaMCPSnorm <- trans_beta$new(dataset = microMCPSnorm, group = "tissue", measure = "bray",ordination = "PCoA")
BetaMCPSnorm

BetaMCPSnorm$plot_ordination(plot_color = "tissue", plot_shape = "Sample_type", plot_group_ellipse = TRUE)
BetaMCPSnorm$cal_manova(cal_manova_set = "tissue*Sample_type")
BetaMCPSnorm$res_manova # Verifying there is no significant interaction between variables


## LEFSe  differential abundance of taxa among tissues. As no singificant interaction exists among tissue and sample type variables
# we evaluated the the differential abundances of ASVs between tissues considering both single and pooled samples in the same group 
LefseMCPSnorm <- trans_diff$new(dataset = microMCPSnorm, method = "lefse", group = "tissue",lefse_norm = 9000 ,alpha = 0.01)
LefseMCPSnorm
LefseMCPSnorm$plot_lefse_bar(LDA_score = 3, keep_prefix = T,simplify_names = F,axis_text_y=10)

LEFSE<-kable(LefseMCPSnorm$res_lefse)
LEFSE # print the formatted results table 


## Functional profiles full dataset

## Functional profiles full dataset
microMCPSnorm
FuncMCPSfull <- trans_func$new(microMCPSnorm)
FuncMCPSfull
FuncMCPSfull$cal_spe_func()
FuncMCPSfull$res_spe_func[1:5,1:30]
FuncMCPSfull$cal_spe_func_perc(use_community = T)
FuncMCPSfull$res_spe_func_perc
FuncMCPSfull$plot_spe_func_perc()  ## default plot of functional profiles

# Extract data and turn it into a matrix
matfuncmcpsFull<-as.matrix(FuncMCPSfull$res_spe_func_perc)
dim(matfuncmcpsFull)
rownames(matfuncmcpsFull)
FuncMCPSfull$sample_table$sampleID
rownames(matfuncmcpsFull)<-FuncMCPSfull$sample_table$sampleID
matfuncmcpsFull

#Transposed matrix for heatmap downstream 
mat2funcmcpsFull<-t(matfuncmcpsFull)
mat2funcmcpsFull
dim(mat2funcmcpsFull)


## Convert matrix into a data frame to generate dotplot and compare means
matfuncmcpsFullDF<-as.data.frame(matfuncmcpsFull)
matfuncmcpsFullDF$tissue<-FuncMCPSfull$sample_table$tissue
matfuncmcpsFullDF


# Melt dataframe
matfuncmcps3FullDF<-melt(matfuncmcpsFullDF)
matfuncmcps3FullDF
matfuncmcps3FullDF$variable <- as.character(matfuncmcps3FullDF$variable) #convert to character

## Data summary function
data_summary <- function(data, varname, groupnames){
  require(plyr)
  summary_func <- function(x, col){
    c(mean = mean(x[[col]], na.rm=TRUE),
      se = se(x[[col]]/sqrt(length(x[[col]])), na.rm=TRUE))
  }
  data_sum<-ddply(data, groupnames, .fun=summary_func,
                  varname)
  data_sum <- rename(data_sum, c("mean" = varname))
  return(data_sum)
}

# Variable containing the data summary (mean by tissue)
dfFunc <- data_summary(matfuncmcps3FullDF, varname="value", 
                       groupnames=c("tissue","variable"))
dfFunc

# Convert categories to a factor variable
dfFunc$tissue=as.factor(dfFunc$tissue)
dfFunc$variable=as.factor(dfFunc$variable)
dfFunc
head(dfFunc)
dfFunc$se
dfFunc$se<-as.integer(dfFunc$se)
dfFunc

# Statistical test assessing differences between tissue types by predicted functions
FUNCstat.test2 <- compare_means(
  value ~ tissue, data = matfuncmcps3FullDF, group.by = "variable",
  method = "wilcox.test", p.adjust.method = "fdr")


SigCat2<-FUNCstat.test2 %>% filter(FUNCstat.test2$p.adj<0.05)
SigCat2

highlight<-filter(dfFunc,variable %in% SigCat2$variable)
highlight  ### DF rows corresponding to the Significantly different Func categories


dotcolor<-c("#1b9e77","#d95f02")


dotplotMCPSFUNC <- ggplot(dfFunc, aes(x= factor (variable, levels = rev (levels(factor(variable)))), y=value, fill=tissue)) + geom_point(aes(color=tissue),size=2.5,alpha=0.2) +coord_flip() + geom_line(aes(group=variable), color="#1E90FF", alpha=0.2) + 
  geom_errorbar(aes(ymin=value-se, ymax=value+se), width=.01, alpha=0.4 )+
  geom_point(data= highlight, aes(color=tissue),size=3.7,alpha=0.8)+geom_line(data=highlight,aes(group=variable), color="#1E90FF", alpha=0.4)+
  geom_line(data= highlight,aes(group=variable), color="#1E90FF", alpha=0.8)+
  geom_errorbar(data= highlight,aes(ymin=value-se, ymax=value+se), width=.08, alpha=0.6 )+
  theme_minimal()+ theme(legend.position="top",axis.text.x = element_text(size=9,face="bold", angle = 0),
                         axis.text.y = element_text(size=10, angle = 0)) +
  scale_color_manual(values = c("crop" = "#1b9e77", "midgut" = "#d95f02")) + labs (x="functional category", y="community percentage capable of executing the predicted functions ")

dotplotMCPSFUNC


# Heatmap created with the pheatmap package

## Side information and annotation colors for heatmap
colnames(FuncMCPSfull$sample_table)
FuncMCPSfull$sample_table[,c(1,2)]

Sideinfo_type_tissue<-FuncMCPSfull$sample_table[,c(1,2)]
rownames(Sideinfo_type_tissue)
rownames(Sideinfo_type_tissue)<-FuncMCPSfull$sample_table$sampleID
Sideinfo_type_tissue<-as.matrix(Sideinfo_type_tissue)
Sideinfo_type_tissue<-as.data.frame(Sideinfo_type_tissue)
Sideinfo_type_tissue


ann_colorsFUNCMCPS=list(tissue=c('crop'="#1b9e77",'midgut'="#d95f02"),Sample_type=c('pool'="black",'single'="#C0C0C0"))
ann_colorsFUNCMCPS


pheatmap(mat2funcmcpsFull,cluster_rows = T,cluster_cols = T,fontsize_col = 9,fontsize_row=9,annotation_col = Sideinfo_type_tissue,annotation_colors =ann_colorsFUNCMCPS ,cellwidth = 20,cellheight = 10,clustering_distance_cols = "manhattan",clustering_method = "ward.D2",clustering_distance_rows = "euclidean",
         color=colorRampPalette(c("white","plum","#9370DB","#483D8B","#4B0082")) (1200)) 


# Verifying the functions that are singletons
## extracting the singletons and ubiquitous categories

countFuncmat<-mat2funcmcpsFull
countFuncmatbinary = ifelse(as.data.frame((countFuncmat))>0,1,0)
countFuncmatbinary
rowSums(countFuncmatbinary)
